# Stabilized ion selectivity corrects activation drift in kalium channelrhodopsins

**DOI:** 10.1101/2025.05.20.654571

**Authors:** Xiao Duan, Chong Zhang, Stanislav Ott, Zhiyi Zhang, Christiane Ruse, Sven Dannhäuser, Robert Jacobi, Nadine Ehmann, Divya Sachidanandan, Georg Nagel, Manfred Heckmann, Robert J. Kittel, Alexander Gottschalk, Adam Claridge-Chang, Shiqiang Gao

## Abstract

Effective optogenetic inhibition of neuronal activity requires tools that can reliably silence neurons across diverse conditions and cell types. Potassium-selective channelrhodopsins (KCRs) have emerged as promising alternatives to chloride-conducting channels for optogenetic inhibition of cellular excitability, but they exhibit dynamic ion selectivity shift under prolonged or intense illumination, causing activation. This inhibition to activation transition limits their utility for silencing circuits. Through behavioral and electrophysiological analyses in *Drosophila*, and *C. elegans*, we found that the C29D variant of KCR1 maintains the most stable potassium selectivity among tested variants. While other KCR variants showed excitation defects, KCR1-C29D consistently provided robust in vivo inhibition across cell types, illumination conditions, and species. Electrophysiological recordings revealed that while most KCRs show declining PK/PNa ratios during illumination, KCR1-C29D maintains stable ion selectivity, consistent with its robust silencing capability. This work resolves a key limitation of KCR optogenetics and shows that KCR1-C29D is a superior tool for reliable inhibition. Our findings offer valuable mechanistic and practical insights for the design of next generation optogenetic inhibitors.

## Introduction

Optogenetic tools like the channelrhodopsins (ChRs) have revolutionized neuroscience by enabling precise control of neuronal activity with light [1–4]. While ChR2 and its variants effectively activate neurons [5, 6], inhibitory optogenetic tools have lagged behind [7]. The halorhodopsins (HRs) like NpHR [8] and eNpHR3.0 [9] are inhibitory chloride pumps that desensitize under constant illumination [10]. Proton pumps such as bacteriorhodopsin (BR) [11, 12] and archaerhodopsin (Arch) [13], hyperpolarize neurons but alter intracellular pH, and can cause rebound firing and calcium influx [14]. All ion pumps suffer from low photon efficiency, require high light intensities, thereby risking phototoxicity and tissue heating.

Anion channelrhodopsins (ACRs) like GtACR1 from *Guillardia theta* [15] offer a more physiological approach by mimicking GABA/glycine signaling. These are highly effective inhibitors, though with some limitations. In mammals, ACRs have been observed to depolarize compartments with high intracellular chloride, potentially causing excitation [14, 16]. In mammalian cells, soma-targeted ACR variants improved inhibition [17]. In *Drosophila*, ACRs have proven widely useful, though broad ACR expression has been linked to toxicity even in the dark [18].

Since potassium (K⁺) channels naturally restore resting membrane potential, K⁺-based optogenetic tools have been pursued for neuronal silencing. Mimicking the natural process of potassium-mediated membrane repolarization to silence neurons has been a longstanding goal in inhibitory optogenetics. Early synthetic light-regulated K⁺ channels included BLINK1 [19] and SthK-bPAC [20, 21], but the latter’s reliance on cAMP led to delayed activation and dark activity, causing eclosion failure in *Drosophila* motor neurons [20]. Despite their limitations, these studies demonstrated that K⁺ channels could effectively silence neurons and highlighted the potential of K⁺-selective optogenetic inhibitors.

The discovery of natural K⁺-selective channelrhodopsins (KCRs), starting with HcKCR1 and HcKCR2 from *Hyphochytrium catenoides*, was a breakthrough [22]. These channels exhibit high K⁺ conductance, enable membrane hyperpolarization. HcKCR1 provided inhibition with minimal phototoxicity and reduced post-illumination rebound [23].

Subsequent work identified another naturally-occurring KCR from *Wobblia lunata*, WiChR [24]; and engineered a highly sensitive HcKCR1 variant, HcKCR1-hs [25]; and mutant KCR variants with improved K^+^ selectivity, e.g. KALI1 [26]. In rodent epilepsy models, HcKCR1-hs inhibited seizure activity and delayed seizure onset via transcranial optogenetic stimulation, underlining KCRs as promising tools for studying neural circuits and treating hyperexcitability disorders.

However, through these studies it became apparent that, under continuous illumination, KCRs can exhibit reduced K^+^ selectivity [22, 24, 26]. Relatively decreased potassium conductance could lead to undesirable neuronal activation. In *Caenorhabditis elegans*, HcKCR1 stimulation in cholinergic neurons induced neuronal inhibition and muscle relaxation under optimized light conditions [27]. Yet, while lower-intensity illumination sustained inhibition in specific transgenic strains, high-intensity continuous illumination caused only transient muscle relaxation. Actuation with WiChR produced similar effects, where prolonged illumination induced depolarization and muscle contraction, likely due to WiChR’s high conductivity and partial conductance of other cations including Na⁺.

These unwanted effects were mitigated by pulsed illumination or less efficient channel activation with green or orange light [27]. Indeed, a 10 ms light pulse delivered at 10 Hz has been used to suppress KA-induced convulsive seizures in mice [25]. While this approach reduces activation events, it is not suitable for achieving long-term, sustained inhibition. Variability may also stem from particular physiological features of different organisms, e.g. membrane composition, ambient temperature, and the repertoire of endogenous ion channels. These observations highlight that KCRs can exert complex effects on neural circuits and are not consistently inhibitory, underscoring the need to systematically investigate the physiological and biophysical factors that influence their performance.

To further investigate the variable effects of KCR actuation, we conducted validation in the vinegar fly *Drosophila melanogaster* and *C. elegans* to compare the inhibitory efficacies of the different KCR variants. We found that, under low-intensity or short-duration illumination, the KCRs mostly inhibited neuronal activity; however, under high-intensity or prolonged illumination many KCR variants became neuronal activators. One exception was KCR1-C29D, which was consistently inhibitory. By analyzing behavioral phenotypes and electrophysiological properties, our study offers valuable insights into the mechanisms underlying KCR inhibition to activation transitions. It also establishes a KCR variant for reliable optogenetic inhibition and provides further guidance for engineering optimized K⁺-selective channelrhodopsins.

## Results

### In *Drosophila,* KCRs can activate neurons

The simplest behavioral experiment to assess KCR function is to monitor the locomotor activity of the animal after ChR actuation in motor neurons. We started our investigations by comparing the inhibitory efficacies of the previously developed KCR1 2.0 variant [28] with GtACR1 [29, 30] (referred to as ACR1 hereafter). As a reference for neuronal activation effects, we used the light-gated cation channel ChR2 carrying the D156H mutation, ChR2-XXM [31] (hereafter referred to as XXM in the figure and) (**Figure 1A**).

**Figure 1:**
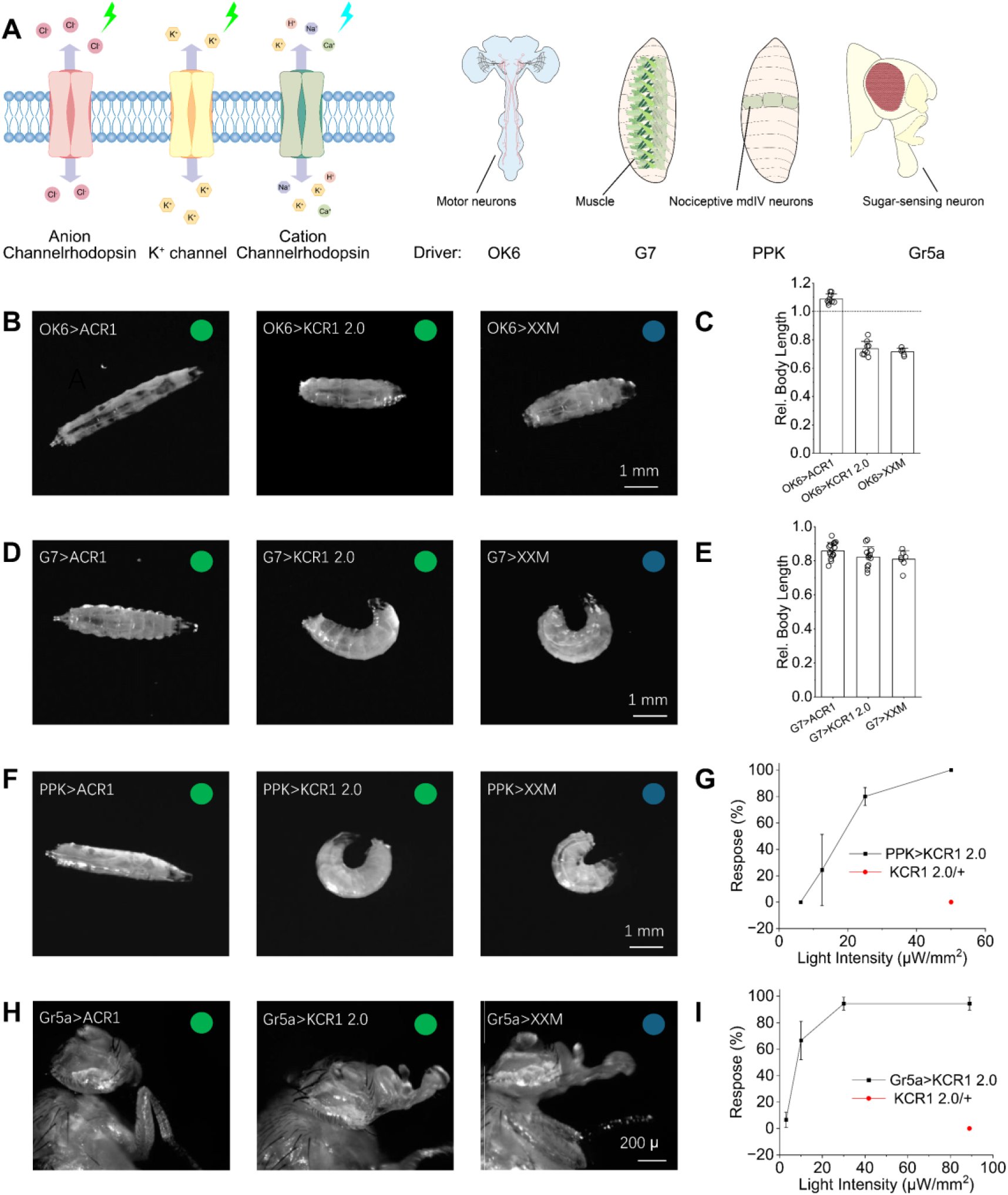
Actuation of KCR1 2.0 activates several *Drosophila* circuits. **A**, Schematic overview of ChRs and GAL4 drivers used in this study. **B**, **D**, **F**, **H**, Representative images showing light-induced body postures in *Drosophila* expressing ACR1, KCR1 2.0, or XXM in motor neurons (**B**), muscle cells (**D**), Class IV multidendritic (mdIV) sensory neurons (**F**) and sugar-sensing neurons (**H**). **C** and **E**, Quantification of larvae body length changes in response to light stimulation when channelrhodopsins were expressed in motor neurons [**C**; mean ± SD, n = 9 (ACR1 and KCR1 2.0), n = 5 (XXM)] and muscle cells [**E**; mean ± SD, n = 15 (ACR1), n = 12 (KCR1 2.0), n = 7 (XXM)]. Relative body length was calculated as the ratio of body length at the end of 20 s illumination epoch compared to the epoch before illumination. In **B** and **C**, larvae were illuminated with green light (530 nm, 28 μW/mm²) for ACR1 and KCR1 2.0, and blue light (450 nm, 313 μW/mm²) for XXM. The same wavelengths were used in **D** and **E**, with adjusted intensities: 80 μW/mm² for ACR1, 19 μW/mm² for KCR1 2.0, and 31 μW/mm² for XXM. In (F), green light (530 nm, 47 μW/mm²) was used for ACR1 and KCR1 2.0, and blue light (473 nm, 63 μW/mm²) for XXM. In (H), green light (530 nm, 73 μW/mm²) was used for ACR1 and KCR1 2.0, and blue light (473 nm, 234 μW/mm²) for XXM. **G**, Percentage of *PPK>KCR1* larvae exhibiting corkscrew-like behavior in response to 10 s illumination at varying light intensities. 15 larvae were considered as one iteration with n = 3 repeats. Data are presented as mean ± SD. **I**, Dependence of PER response rate on light intensity in *Gr5a>KCR1* flies. Illumination time was 20 s. A set of 15 flies was regarded as one iteration, with n = 3 repeats. Data are presented as mean ± SD.

With the OK6-GAL4 driver line, ACR1 actuation in motor neurons induced body relaxation, whereas XXM actuation caused body contraction (**Figure 1B, C**). KCR1 2.0 activation initially led to a brief relaxation phase, followed by contraction. This suggested that prolonged illumination induced a transition from neuronal inhibition to activation. In muscle cells by *G7-GAL4*, both KCR1 2.0 and XXM induced body contraction and curling, whereas ACR1 also led to body contraction (**Figure 1D, E**).

We next expressed ChRs in Class IV multidendritic (mdIV) sensory neurons using the *PPK-GAL4* driver line. Activation of these neurons triggers nociceptive behavioral responses, most notably the “corkscrew” body roll with a rapid rotation along the animal’s long axis [32, 33]. Activation of KCR1 2.0 induced corkscrew-like body rolling, resembling the behavioral response elicited by XXM (**Figure 1F, G**). The proportion of *PPK>KCR1 2.0* larvae exhibiting nociceptive responses was positively correlated with light intensity, while no such response was observed in control and ACR1 larvae. All *PPK>KCR1 2.0* larvae initiated rolling responses upon exposure to 50 µW/mm² green light. However, the rolling response occurred almost immediately upon illumination in *PPK>XXM* larvae, whereas the responses of *PPK>KCR1 2.0* larvae occurred several seconds after illumination. This delay aligns with the gradually decreasing potassium selectivity of KCRs [22, 24]. To verify that neurons were silenced by ACR1 actuation, we applied a mechanical stimulus to the larval posterior region (around segment A8) with a 2840 kPa pressure filament during illumination. ACR1 effectively suppressed the nocifensive response, demonstrating its inhibitory effect on nocifensive signaling (**Supplemental Figure 1**).

We next expressed the three ChRs in sugar-sensing gustatory neurons using the *Gr5a-GAL4* driver. Previous studies have shown that the proboscis extension response (PER) can be triggered by activation of ChR2 and its variants [34]. We observed that KCR1 2.0 also elicited the PER upon green light stimulation (**Figure 1H, I**). As light intensity increased from 3 µW/mm² to 30 µW/mm², a higher proportion of flies exhibited proboscis extension during illumination. At 30 µW/mm², up to 94% of flies displayed PER, with no further increase noted at ∼90 µW/mm². In contrast, no PER was detected in *Gr5a>ACR1* flies under any illumination conditions.

These experiments demonstrate that KCR1 2.0 unexpectedly activates multiple fly circuits. While initially providing brief inhibition, prolonged illumination causes KCR1 2.0 to transition to activation, producing contraction via motor neurons, nociceptive responses via sensory neurons, and proboscis extension after actuation of gustatory neurons. This inhibition to activation shift in vivo highlights the need for KCRs with straightforward inhibitory effects.

### KCR1-C29D consistently silences neurons in *Drosophila* larvae

The activation-associated phenotypes observed with KCR1 2.0 prompted us to re-examine several of the previously published KCR constructs. We first analyzed *Drosophila* larvae in which opsin was expressed was induced in motor neurons using the *OK371-GAL4* driver. As shown in **Supplemental Figure 2**, motor neuron activation with the cation channel *CsChr* as well as inhibition with ACR1 produced a reduction in speed, indicating that locomotion suppression alone can not be used as a reliable behavioral marker for distinguishing between neuronal activation and silencing.

To identify clear differences between CsChr (activation), ACR (inhibition), and KCR actuation phenotypes, we therefore analyzed distinct behavioral states in detail so as to reliably discriminate neuronal excitation from silencing. Targeting motor neurons with *OK371>CsChr* induced immobility with whole body contraction (**Figure 2A,B**), similar to activation behavior in *C. elegans* [27, 35]. By contrast, actuating the neurons with *OK371>ACR1* resulted in immobility without contraction, consistent with neuronal silencing (**Figure 2A,C**). Actuation with KCR1 2.0, WiChR and KCR1-ET produced contraction phenotypes similar to CsChr, whereas actuation with KCR1-C29D resulted in immobility without contraction, as observed with ACR1 (**Figure 2D-F**). We also tested flies that expressed the recently developed HcKCR KALI (H225F) variants [26].

**Figure 2:**
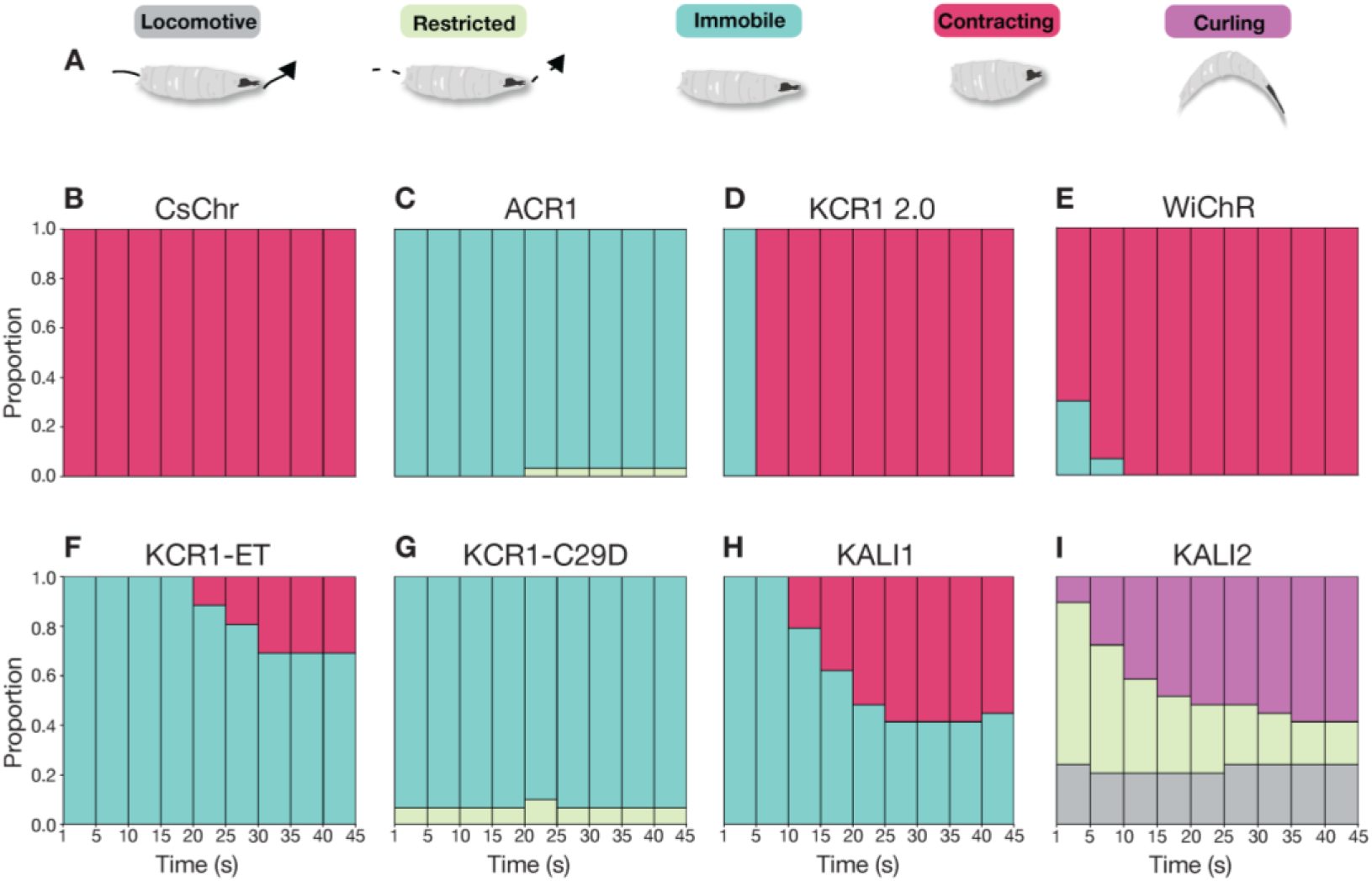
Motor neuron actuation with KCR variants induces different behavioral states in *Drosophila* larvae. **A**, Schematic and colour legend of behavioral states in *Drosophila* larvae induced by opsin actuation. **B**, State bar chart during light illumination for *Drosophila* larvae expressing *OK371>CsChr*. The illumination epoch (45 s) was divided into 5 s bins and the prevalence of each state across animals was plotted as a proportion in each bin (see methods). Activation of motor neurons with CsChr induced immobility and body contraction in all animals. **C**, ACR1 actuation resulted in immobility without body contraction throughout the illumination epoch. **D**, KCR1 2.0 activation induced body contraction approximately 5 s into the illumination epoch. **E**, WiChR activation induced body contraction in the majority of larvae from the start of light exposure. **F**, Actuation with KCR1-ET predominantly led to immobility, with some contraction phenotypes observed at the later stages of illumination. **G**, Silencing with KCR1-C29D induced immobility, similar to the phenotype observed with ACR1. **H**, KALI1 actuation produced a mixed phenotype characterized by immobility and partial body contraction at later illumination stages. **I**, KALI2 activation induced curling and led to incomplete locomotor impairment. Red light: λ = 617 nm, 13 μW/mm^2^; Green light: λ = 530 nm, 22 μW/mm^2^; Blue light λ = 460 nm, 21 μW/mm^2^. Sample size: *OK371>CsChr* n = 29 larvae, *OK371>ACR* n = 30 larvae, *OK371>KCR1 2.0* n = 29 larvae, *OK371>WiChR* n = 30 larvae, *OK371>KCR1-ET* n = 26 larvae, *OK371>KCR1-C29D* n = 30 larvae, *OK371>KALI1* n = 29 larvae, *OK371>KALI2* n = 29 larvae.

Compared to wild type HcKCRs, the KALI variants were reported to display improved K^+^ selectivity and superior inhibitory properties. However, actuated *OK371>KALI1* larvae displayed partial contraction while actuation of KALI2 in motor neurons produced curling, and many larvae remained locomotive (**Figure 2 G,H**). Comparable results were obtained by using pan-neuronal *elav-GAL4* (**Supplemental Figure 3A-E**) and the motor neuron driver *OK6-GAL4* (**Supplemental Figure 3F-I**), suggesting that the observed effects are generalizable to different neuronal subsets. Taken together, this analysis showed that five recently described KCR variants (KCR1 2.0, WiChR, KCR1-ET, KALI1, and KALI2) all elicited responses similar to a non-selective cation channel. This analysis also established that KCR1-C29D elicited effects similar to an anion channel, and is therefore likely the most effective KCR variant for neuronal inhibition in flies.

To further compare the inhibitory effects of different KCR variants, we monitored their locomotor responses following short-term stimulation (< 3 s) while gradually increasing light intensity (**Figure 3**). In this experiment, we also tested a novel WiChR T95C mutant, which exhibited a slightly negative reversal potential and faster kinetics (**Supplemental Figure 4**), as well as a blue-shifted KCR1-T136A/G140A mutant (KCR1-TAGA) [26]. Additionally, we generated soma-targeted versions of WiChR-T95C (sWiChR-T95C) and KCR1-TAGA (sKCR1-TAGA), mimicking the soma-targeted ACR1 [17].

**Figure 3.**
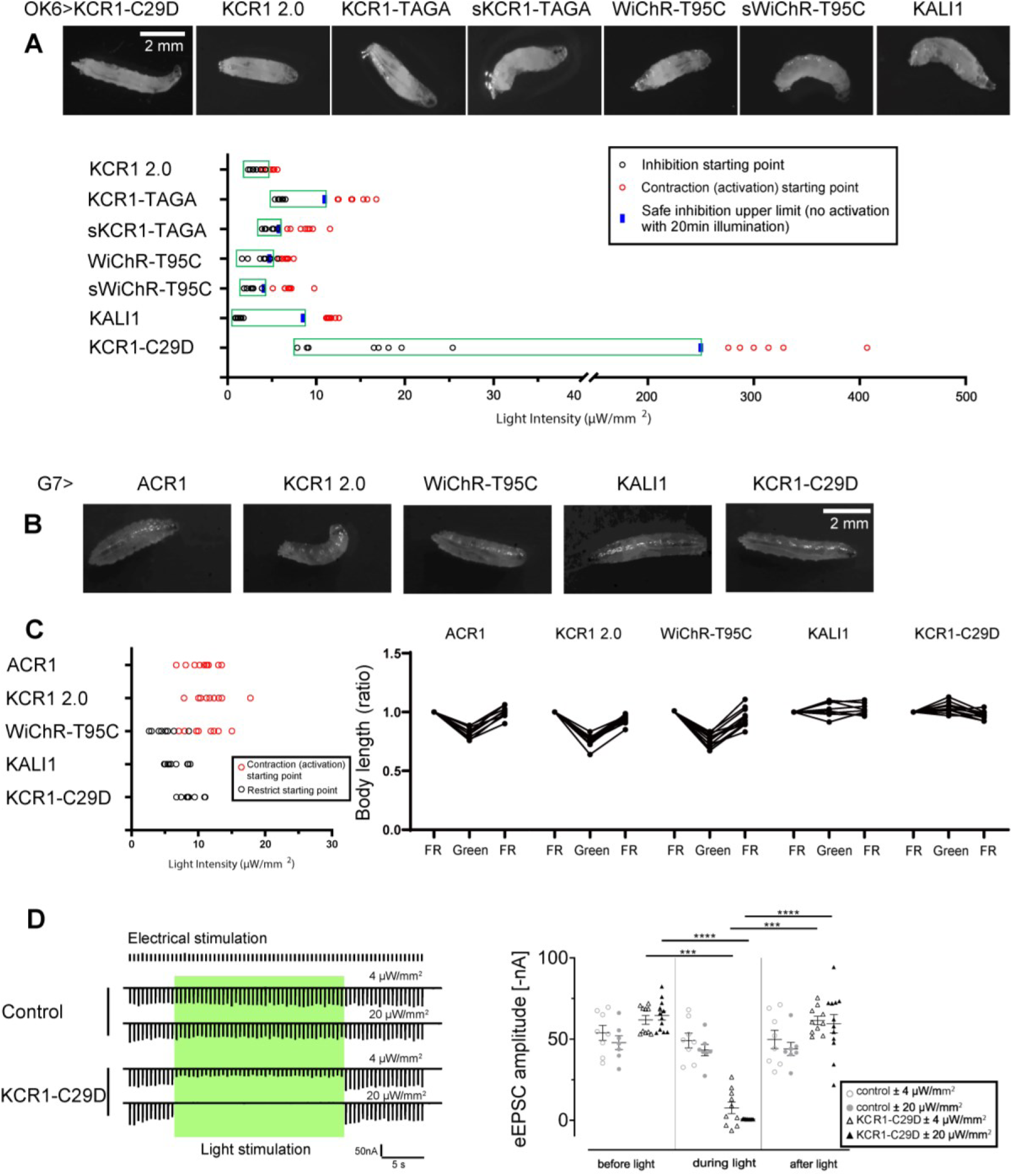
Evaluation of inhibitory and activating effects of KCRs in motor neurons and muscle cells. **A**, Representative images from video recordings of third-instar larvae expressing KCR1 2.0, KCR1-TAGA, sKCR1-TAGA, KALI1, KCR1-C29D, WiChR-T95C, and sWiChR-T95C in motor neurons (*OK6-GAL4*) following green light illumination (λ = 530 nm, light intensity: 20 µW/mm²). Behavioral responses were tested by gradually increasing light intensity. The safe inhibition limit was defined as the maximum light intensity below the activation threshold, confirmed by maintaining inhibition during 20 minutes of continuous illumination. n = 6 - 10 per group, all data points were shown. **B**, Representative images from video recordings of third-instar larvae expressing different KCR variants in muscle cells (*G7-GAL4*) under green light illumination (λ = 530 nm, light intensity: 20 µW/mm²). **C**, Behavioral responses of larvae expressing KCR variants in muscle cells across different light intensities. The restricted (immobilized) state was not observed in ACR1 and KCR1 2.0, whereas the contracted state was not observed in KALI1 and KCR1-C29D. n = 8 - 10, all data points were shown. **D**, Electrophysiological analysis of light-induced inhibition of eEPSCs (evoked excitatory postsynaptic currents) at the neuromuscular junction (NMJ) comparing control and *OK6>KCR1-C29D* larvae, green light illumination condition: λ = 530 nm; light intensity, 4 and 20 µW/mm², n = 7 - 12, all data points were shown, error bars = SD.

However, our imaging analyses showed that soma-targeting signals derived from mammalian neurons did not function well in *Drosophila*, and no apparent increase in soma-targeted expression was observed in comparison to other KCRs (**Supplemental Figure 5 A-F**). Both soma-targeted KCRs also exhibited a transition from inhibition to activation over time under illumination of 21 µW/mm², 460 nm light (**Supplemental Figure 5 G,H**), similar to KCR1 2.0 and WiChR shown in **Figure 2D-E**, albeit to a lesser extent.

As shown in **Figure 3A**, most KCR variants exhibited a very narrow transition phase from inhibition to activation when shifting from lower to higher light intensity. Among them, KCR1-TAGA, KALI-1, and KCR1-C29D demonstrated a relatively broader inhibition range. Notably, KCR1-C29D exhibited a convincing inhibitory effect across a light intensity range of 10 µW/mm² to approximately 250 µW/mm². Furthermore, we confirmed that illumination at 250 µW/mm² could suppress larval locomotion for at least 20 minutes with KCR1-C29D. At around 300 µW/mm², KCR1-C29D triggered body contraction, but prolonged global illumination at this intensity resulted in lethality after 30 minutes. Therefore, such high-intensity illumination should be avoided in experimental settings. We then tested the behavioral responses to stepped increases in light intensity following expression of several KCRs in the muscles. Consistent with previous observations, KCR1-C29D exhibited an inhibitory effect at low light intensity (10 µW/mm²) without a clearly defined activation phase. Similarly, muscle expression of KALI induced inhibition at 10 µW/mm² without triggering obvious activation (**Figure 3B**). Notably, although expression of these KCRs effectively suppressed larval body movement without causing contraction under illumination, head movements remained detectable. Measurements of larval body length further corroborated these findings, corresponding to either the inhibitory or contractile states observed (**Figure 3C**).

These data highlighted KCR1-C29D as the most promising optogenetic inhibitor for *Drosophila* larvae among the currently available KCR variants. Electrophysiological recordings at the neuromuscular junction (NMJ) further demonstrated that evoked excitatory postsynaptic currents (eEPSCs) can be effectively inhibited by KCR1-C29D. A low light intensity of 4 μW/mm² significantly reduced eEPSC amplitudes, while a higher intensity of 20 μW/mm² completely abolished them (**Figure 3D; Supplemental Tables 1 and 2**).

### KCR1-C29D, KCR1-ET and KALI1 silence motor neurons in *Drosophila* adults

To investigate whether the activation-associated phenotypes induced by KCR are also present in adult *Drosophila*, we expressed CsChr, ACR1 and KCRs in motor neurons (*OK371-GAL4*) and assessed fly posture in a horizontal locomotion assay [18]. As observed in larvae (**Supplemental Figure 1**), actuation of all opsins besides KALI2 also impaired locomotor activity in adult flies (**Supplemental Figure 7**). However, a closer analysis of fly posture during light stimulation revealed differences between CsChr, ACR1 and KCR actuation (**Figure 4**). Activation of motor neurons with CsChr induced a unique phenotype that was characterised by spasms and a transition from an angled to a supine posture throughout the illumination period (**Figure 4A,B**). Silencing with ACR1 induced immobility and a supine fly posture without limb movement (**Figure 4A,C**), whereas silencing with KCR1-ET, KCR1-C29D and KALI1 led to a prone posture where some restricted walking occurred (**Figure 4A, D-E, G**). Motor neuron actuation with other KCR variants induced supine postures and abdomen curling that was especially pronounced in flies expressing the WiChR variants (**Figure 4F, G, J-K**). Taken together, these data indicate that optogenetic actuation of motor neurons by WiChR, KCR2.0 and their respective somatic variants induce postural changes that are associated with muscle contraction and potential neuronal activation [27, 35] in *Drosophila* adults as well as in larvae. By contrast, actuation with KCR1-C29D, KCR1-ET and KALI1 do not elicit neuronal firing. However, their silencing efficacy appears to be somewhat weaker when compared to ACR1 at the light intensity of 15 μW/mm^2^.

**Figure 4:**
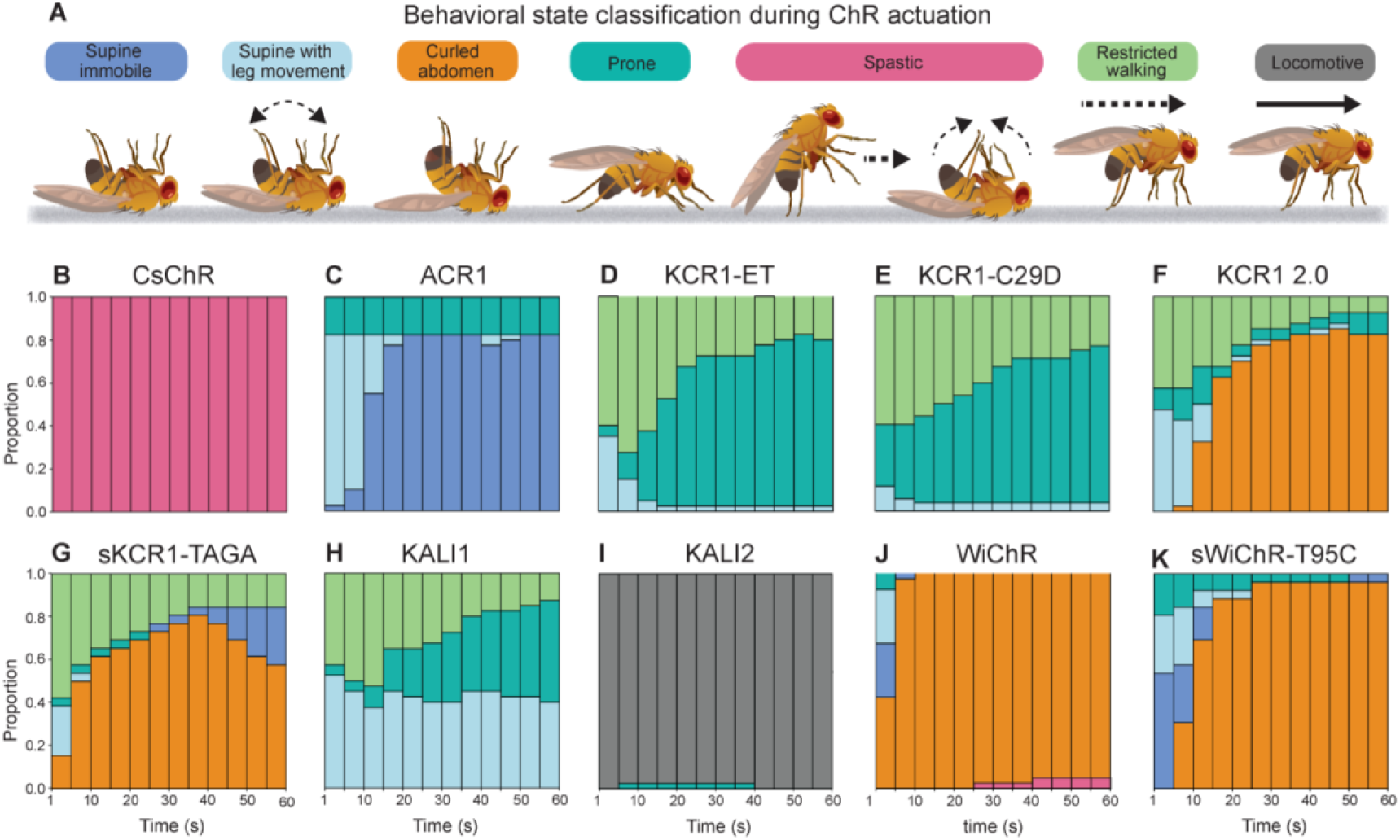
Motor neuron actuation with CsChr, ACR and KCRs induces different behavioral states in adult *Drosophila*. **A**, Schematic representation and colour legend of behavioral states in *Drosophila* adults induced by respective opsin activation. **B**, CsChr activation produced a distinct spastic phenotype characterized by cramped muscles and a rigid stature. The fly posture transitioned from inclined to supine with leg folding throughout the illumination period. **C**, ACR1 activation elicited the strongest inhibitory phenotype that led to a supine or prone posture. **D-E**, KCR1-ET and KCR1-C29D activation impaired locomotion and induced a prone posture, with occasional restricted walking. **F-G**, Activation of KCR1 2.0 and sKCR1-TAGA produced a supine posture and abdomen curling as well as occasional restricted walking. **H-I**, KALI 1 actuation induced a heterogeneous phenotype that consisted of supine and prone postures as well as restricted walking (**H**). KALI2 actuation was largely ineffective and did not impair locomotion (**I**). **J-K**, Activation of WiChR and sWiChR-T95C induced a supine posture and abdomen curling. In all panels the opsins were expressed in motor neurons with *OK371-GAL4*. Genotypic driver and responder controls were used in each experiment. Red light: λ = 617 nm, 14 μW/mm^2^; Green light: λ = 530 nm, 15 μW/mm^2^; Blue light λ = 460 nm, 18 μW/mm^2^. Sample size: *OK371>CsChr* n = 52, *OK371>ACR1* n = 40, *OK371>KCR1-ET* n = 40, *OK371>KCR1-C29D* n = 52, *OK371>KCR1 2.0* n = 40, *OK371>sKCR1-TAGA* n = 26, *OK371>KALI1* n = 40, *OK371>KALI2* n = 40, *OK371>WiChR* n = 40, *OK371>sWiChR-T95C* n = 26.

### KCR1-C29D actuation does not activate gustatory neurons

We further validated the silencing efficiency of KCR1-C29D and the newly generated KCR variants by conducting additional PER and memory induction experiments. To investigate whether KCR actuation would lead to neuronal activation at higher light intensities we expressed the different opsins in sugar-sensing neurons with *Gr5a-GAL4* and exposed the flies to a light ramp. The majority of KCRs displayed PER at varying light intensities, with somatic KCRs and WiChR requiring the least amount of light to elicit PER (**Figure 5A,B**). Although 50% and 30% of KALI1- and KALI2-expressing flies did indeed display PER at comparatively higher light intensities, the rest of the tested KALI population did not show PER at intensities of up to 320 μW/mm^2^. By contrast, neither ACR1- nor KCR1-C29D-expressing flies displayed PER, suggesting that actuation of these opsins does not activate neurons.

**Figure 5:**
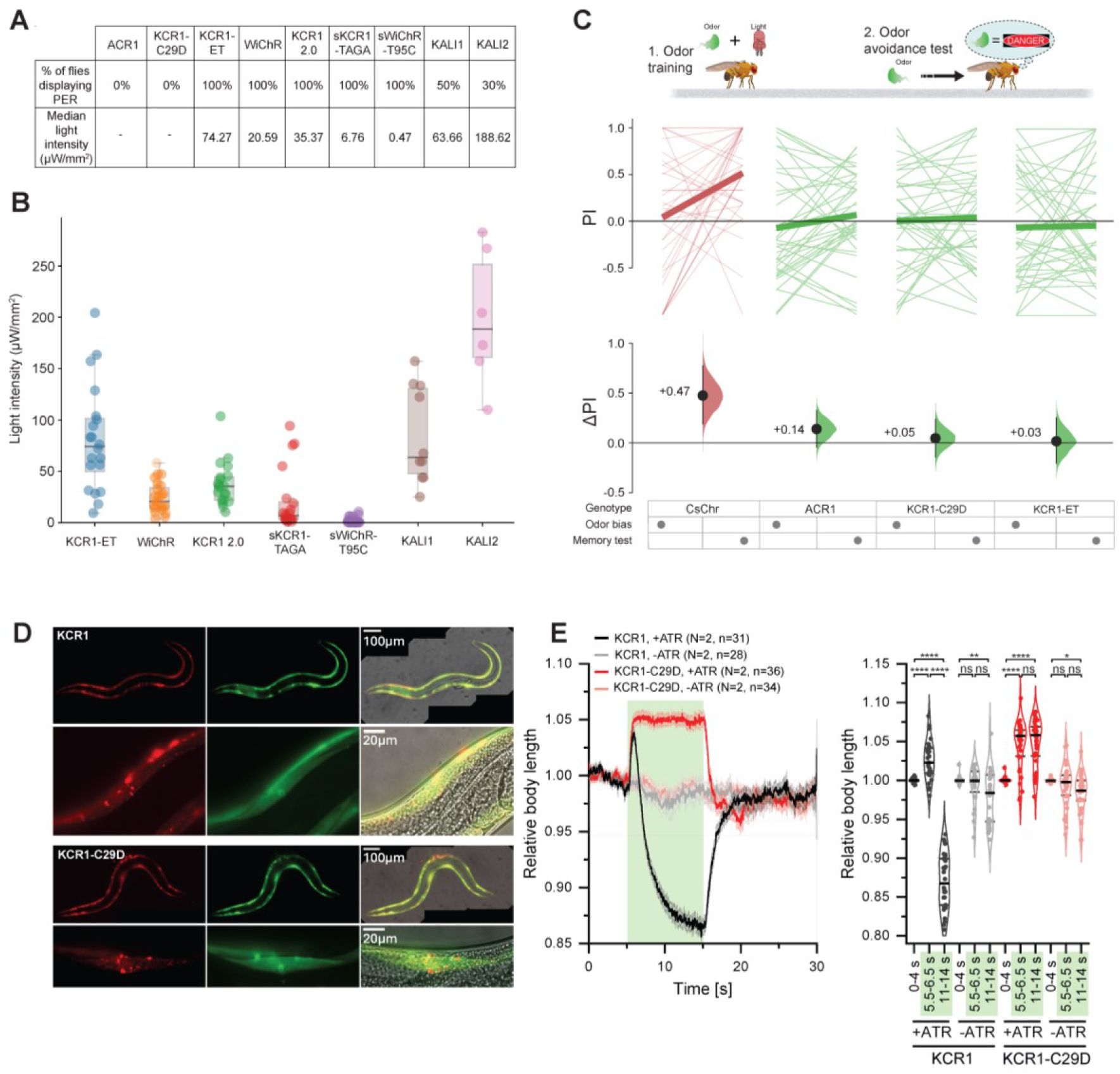
Neuronal actuation with KCR1-C29D silences *Drosophila* gustatory and memory neurons and *C. elegans* muscles. **A**, A summary table of % flies displaying PER and corresponding median light intensities for the respective genotypes. Opsins were expressed in gustatory neurons by using *Gr5a-GAL4*. **B**, A box plot displaying the median and individual PER responses of *Drosophila* at the respective light intensities. Each dot represents one fly. Error bars show 95% CI. Sample size: *Gr5a>KCR1-ET*: n = 20 flies, *Gr5a>WiChR*: n = 24 flies, *Gr5a>KCR1 2.0*: n = 20 flies, *Gr5a>sKCR1-TAGA*: n = 20 flies, *Gr5a>sWiChR-T95C*: n = 25 flies, *Gr5a>KALI1*: n = 20 flies, *Gr5a>KALI2*: n = 20 flies. **C**, Pairing odor stimulation with the activation of PPL1 neurons by *MB320C>CsChr* induced aversive odor memory. Memory formation did not occur when PPL1 neurons were actuated by KCR1-C29D or KCR1-ET. The schematic above illustrates the experimental setup. The top plot shows *Drosophila* odor avoidance PI during the unpaired test (“bias”, left side) and the odor combined with PPL1 stimulation test (“memory test”, right side). The bottom plot illustrates the mean difference effect sizes between the bias and memory test for each genotype. Sample size: *MB320C>CsChr* n = 210 flies; *MB320C>ACR1* = 228 flies; *MB320C>KCR1-C29D* = 240 flies; *MB320C>KCR1-ET* = 228 flies. Error bands show 95% CI. **D**, Representative images showing the expression of KCR1 (WT) and KCR1-C29D (each as a C-terminal fusion with mCherry; green fluorescence is GFP, expressed from an SL2-linked mRNA) in the body wall muscle cells of *C. elegans*. **E**, Behavioral profiles of *C. elegans* transgenic strains expressing KCR1 (WT) or KCR1-C29D (*pmyo-3::KCR1(WT or C29D)::mCherry::SL2::GFP*, 15 ng/µL) under conditions with and without additional all-*trans* retinal supplementation, illuminated for 10 s with 535 nm light at an intensity of 1mW/mm^2^. Statistical significance was assessed using a mixed-effect model analysis model (REML) with Tukey’s multiple comparisons test.

We next expressed selected opsins in a dopaminergic cluster of PPL1 neurons by using *MB320C-GAL4* to validate KCR1-C29D-mediated silencing in an additional behavioral assay that is able to differentiate between neuronal activation and inhibition. Previous studies have shown that activation of PPL1 neurons elicits aversive memory formation [36]. Accordingly, we observed that pairing odor stimulation with PPL1 activation by CsChr induces conditioned odor avoidance (ΔPI = +0.47, **Figure 5C**) in an olfactory conditioning paradigm [37]. Actuation of PPL1 neurons by ACR1 produced a mild aversive effect (ΔPerformance Index (PI) = +0.14), indicating that ACR stimulation could potentially induce transient neuronal firing or post-inhibitory rebound activity. By contrast, PPL1 actuation by KCR1-C29D or KCR1-ET did not induce odor avoidance (ΔPI = +0.05 and ΔPI = +0.03 respectively), suggesting that these KCR variants do not elicit neuronal firing.

### KCR1-C29D can effectively silence *C. elegans* body wall muscle activity

To extend our *Drosophila* KCR1-C29D findings towards other models, we compared the functionality of KCR1 and KCR1-C29D in body wall muscle cells of *Caenorhabditis elegans*. We previously showed that KCR1 can hyperpolarize *C. elegans* cells only briefly, likely because of a dissipation of the K^+^ gradient, before the Na^+^ current takes over and induces depolarization [27]. Both KCR1 and KCR1-C29D showed similar expression patterns (**Figure 5D**). KCR1-C29D actuation induced stable muscle relaxation, evident as body elongation (corresponding to hyperpolarization) and complete cessation of movement upon green light stimulation. By contrast, KCR1 actuation led to only a brief elongation followed by persistent contraction (corresponding to depolarization); the latter manifesting as egg laying (**Figure 5E** and **Supplemental Figure 7**). Notably, locomotion in KCR1-C29D-expressing animals resumed rapidly after light offset, in contrast to the prolonged recovery period seen with wild-type KCR1 (**Supplemental Video 1**), likely reflecting better preservation of the potassium gradient during illumination. These findings were consistent in two additional independent transgenic lines for each KCR1 and KCR1-C29D (**Supplemental Figure 7**), highlighting KCR1-C29D as a significantly more reliable and efficient optogenetic inhibitory tool in *C. elegans*, compared to WT KCRs tested previously [27].

### The KCR variant KALI1 silences *Drosophila* mechanosensory and memory neurons

We extended the validation of KCR1-C29D and KALI1 in *Drosophila* by expressing the opsins with R18G08-GAL4 in a subset of somatosensory neurons that are involved in the integration of touch stimuli [38]. Activity in similar neuronal subsets has been previously associated with locomotor induction [39] and actuating *R18G08* neurons with CsChr elicited a locomotor response upon light exposure (**Supplementary Figure 8A**). By contrast, actuation with ACR1, KCR1-C29D and KALI did not produce a locomotor response and substantially reduced *Drosophila* activity after the flies have been subjected to an air puff agitation stimulus (**Supplementary Figure 8B-D**). The reduction in locomotion after air puff agitation was similar between ACR1 and KCR variants, indicating that KALI, as well as KCR1-C29D can potently silence fly somatosensory neurons.

To further assess the silencing efficiency of newly generated KCRs in higher order *Drosophila* neurons, we expressed them with *MB247-GAL4* in mushroom body Kenyon cells that are required for associative olfactory memory formation [40]. Flies expressing KCR variants were tested for aversive odor memory formation in a Pavlovian conditioning paradigm by using a multifly olfactory trainer [37]. Actuation of KALI1 and sKCR1-TAGA strongly impaired conditioned odor avoidance (ΔPI = -0.36 and ΔPI = - 0.36 respectively, **Supplemental Figure 9A,C**) that was restored upon retest in the absence of light stimulation. The inhibition of odor memory by other KCR variants was less effective (**Supplemental Figure 9B,D,E**), suggesting that besides KCR1-C29D, the KALI1 variant could serve as an alternative KCR for potent neuronal inhibition of higher-order brain functions.

### Electrophysiological characteristics of KCRs that underlie activation and inhibition

The varying effects of different KCRs on neuronal activity provide a basis for understanding their distinct characteristics. To investigate the mechanisms behind this variation, we conducted electrophysiological studies to characterize representative KCRs, aiming to identify the key factors that govern neuronal inhibition and activation.

We selected KCR1 2.0, WiChR, and WiChR-T95C because they could easily trigger neuronal activation in *Drosophila*, while KCR1-C29D and KALI1 were chosen as representative variants with stronger inhibitory output. Compared to KCR1 2.0, the C29D mutation exhibited a more negative reversal potential and a higher P_K_/P_Na_ (K^+^ permeability to Na^+^ permeability) ratio (**Figure 6A,B and Supplemental Figure 10**), consistent with previous findings [18] and aligned with its stronger inhibitory effect (**Figure 3A**). However, the improved inhibition with KCR1-C29D cannot simply be explained by the enhanced P_K_/P_Na_ ratio alone, since both WiChR and WiChR-T95C showed even more negative reversal potentials and higher P_K_/P_Na_ ratios (**Figure 6C,D and Supplemental Figure 11**).

**Figure 6.**
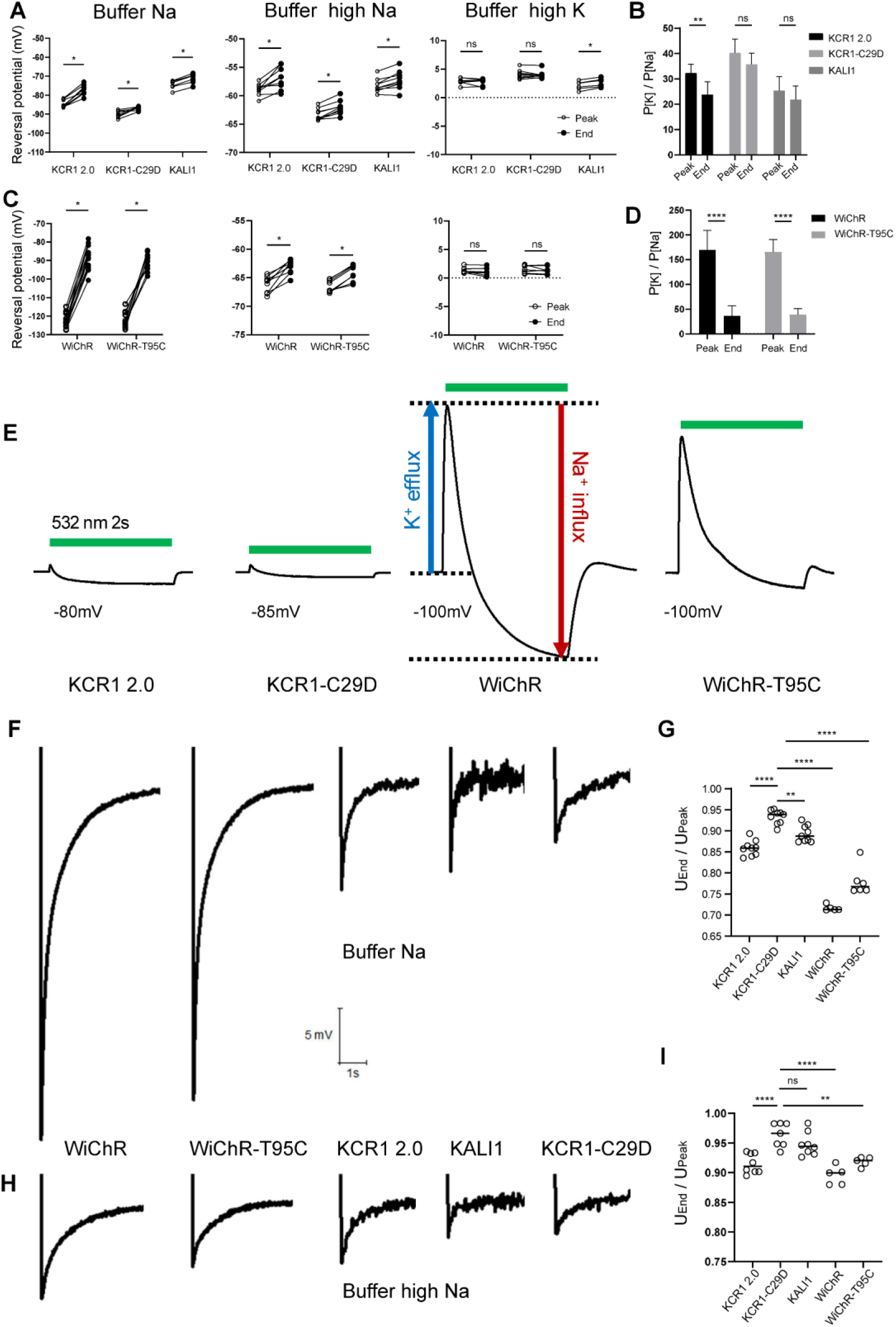
Electrophysiological comparison of KCR variants. **A**, Reversal potentials measured at the peak response and at the end of a 2 s light pulse for KCR1 2.0, KCR1-C29D, and KALI1 in different extracellular buffers. Buffer Na (110 mM NaCl, 2 mM BaCl₂, 1 mM MgCl₂, 5 mM HEPES), Buffer high Na (110 mM NaCl, 5 mM KCl, 2 mM BaCl₂, 1 mM MgCl₂, 5 mM HEPES), and Buffer high K (110 mM KCl, 5 mM NaCl, 2 mM BaCl₂, 1 mM MgCl₂, 5 mM HEPES), all adjusted to pH 7.4. All data points are shown, n = 6 - 9, statistical significance was assessed using multiple paired t-tests. **B**, The K⁺/Na⁺ permeability ratios (PK/PNa) for KCR1 2.0, KCR1-C29D, and KALI1 were determined at the peak response and at the end of the 2 s light pulse. Permeability ratios were calculated using the modified Goldman-Hodgkin-Katz equation, based on the reversal potentials measured in Buffer Na and Buffer high Na. n = 6 - 9, data is shown as Mean ± SD. **C**, Reversal potentials measured at the peak response and at the end of a 2 s light pulse for WiChR and WiChR-T95C in different extracellular buffers. All data points are shown, n = 6 - 9, statistical significance was assessed using multiple paired t-tests. **D**, The PK/PNa ratios for WiChR and WiChR-T95C were determined at the peak response and at the end of the 2 s light pulse. n = 8, data is shown as Mean ± SD. **E**, Representative photocurrent traces of KCR1 2.0, KCR1-C29D, WiChR and WiChR-T95C near their respective reversal potentials. The green bar indicates the period of light illumination. **F**, Representative voltage recording traces of KCR1 2.0, KCR1-C29D, WiChR and WiChR-T95C during light illumination in buffer Na. **G**, Ratios to show the voltage change from peak to the end (2 s after the peak) in buffer Na. All data points are shown, n = 5 - 9. **H**, Representative voltage recording traces of KCR1 2.0, KCR1-C29D, WiChR and WiChR-T95C during light illumination in buffer high Na. **I**, Ratios to show the voltage change from peak to the End (2 s after the peak) in buffer high Na. All data points are shown, n = 5 - 8.

Another key difference is the change in P_K_/P_Na_ ratio during illumination. Both KCR1 and WiChR have been reported to exhibit a decrease in P_K_/P_Na_ ratio upon light activation [22]. The C29D mutant showed no significant change in P_K_/P_Na_ ratio between the peak and end of illumination. Similarly, KALI1 did not show a significantly enhanced P_K_/P_Na_ ratio compared to KCR1 2.0, but like C29D, its ion selectivity remained relatively stable throughout illumination, particularly in buffer close to physiological conditions (e.g., 110 mM NaCl / 5 mM KCl). This suggests that C29D and H225F mutations stabilize the channel’s selectivity, slowing the illumination-induced decrease in P_K_/P_Na_ ratio.

By contrast, WiChR and KCR1 2.0 exhibited a marked decrease in P_K_/P_Na_ during illumination, leading to a transient increase in Na⁺ influx (**Figure 6E**). This transient Na⁺ entry induced a brief depolarization (**Figure 6F-I**), which may trigger action potentials [30, 31]. In the case of WiChR, despite its higher P_K_/P_Na_ and more negative reversal potential, the rapid shift in selectivity causes a stronger Na⁺-driven depolarizing spike (**Figure 6F, H**), capable of triggering action potentials, an effect reminiscent of transient depolarizations observed following hyperpolarization in ventricular myocytes [41].

Moreover, the initial hyperpolarization to more negative potentials (e.g., -100 mV) may relieve the inactivation of voltage-gated Na⁺ channels, enhancing subsequent action potential initiation and neurotransmitter release [42], and likely contributes to the excitatory output observed with WiChR.

## Discussion

In this study, we systematically evaluated the efficacies of various KCRs in silencing activity in *D. melanogaster* and *C. elegans*. Our investigations provide important mechanistic insights and highlight the KCR1-C29D variant as a promising optogenetic inhibitor. While KCRs were originally introduced as potent tools for neuronal silencing due to their high potassium selectivity and low phototoxicity, our results reveal that their inhibitory function can be compromised under prolonged or intense illumination, sometimes resulting in neuronal activation. Consistent with the recent report from *C. elegans* [27], most KCRs tested in this study, including KCR1 2.0 and WiChR, produced transient inhibition followed by activation when exposed to continuous illumination.

Behaviors associated with neuronal activation were consistently observed across different paradigms and model organisms. In *Drosophila* larvae and *C. elegans*, actuation of most KCR variants led to body contraction (**Figures 2,3 and 5E**), whereas KCR activity in *Drosophila* adults elicited the PER and curled abdomen postures (**Figures 4 and 5A-B**). This likely results from light-induced changes in ion selectivity, where initial high K⁺ permeability gradually gives way to increased Na⁺ conductance, thereby leading to a transient depolarization. The KCR1-mediated dissipation of the cellular K^+^ gradient seized K^+^ currents, while the electrochemical driving force for Na^+^ influx persists [27]. Electrophysiological recordings confirmed this shift, revealing a dynamic decrease in the P_K_/P_Na_ ratio during illumination for KCR1 2.0 and WiChR, resulting in net inward Na⁺ currents that may trigger action potentials.

In contrast, the KCR1-C29D mutant displayed a more stable ion selectivity profile, maintaining a relatively constant P_K_/P_Na_ ratio throughout light exposure. Combined with its more negative reversal potential, this stability likely underlies the robust and sustained inhibition observed across a broad range of light intensities and durations.

This preservation of ion selectivity appears to be a critical factor contributing to the superior inhibitory performance of KCR1-C29D. Similarly, KALI1, the second-best KCR variant in our study, showed a stable but not much higher P_K_/P_Na_ ratio during illumination. These findings suggest that a reliable potassium-based optogenetic inhibitor should ideally possess both a stable and relatively high P_K_/P_Na_ ratio. While previous studies have emphasized a high P_K_/P_Na_ ratio as essential for an ideal K⁺-based inhibitory tool, our data highlight that the stability of the P_K_/P_Na_ ratio is more important, provided that the ratio reaches an optimal range.

Moreover, the P_K_/P_Na_ ratio of KCRs seemed to be influenced by multiple factors beyond just the light activation profile. While continuous light illumination activates the channel and impacts ion selectivity, our experiments revealed that buffer composition also significantly affects the P_K_/P_Na_ ratio. As indicated by the reversal potential, the P_K_/P_Na_ ratio is substantially lower in high K^+^ buffer compared to high Na^+^ buffer. It can also be calculated from previous studies that the P_K_/P_Na_ ratio of KCR changed from about 40 to 2, when changing from high Na^+^ to high K^+^ buffer, based on the previously published data from Tajima et al. (**Figure 6A** in [26]). The KCR1-C29D variant, with its stable and relatively high selectivity, represents a robust tool for optogenetic silencing in *Drosophila*. The P_K_/P_Na_ ratio calculated from high K^+^ buffer is also obviously lower for KCR1 than the ratio from the high Na^+^ buffer, which is less significant for KALI1 and not significant for KCR1-C29D (**Supplemental Figure 12**), indicating that unstable P_K_/P_Na_ ratio is influenced by factors such as ionic environment or voltage. It is also conceivable that KCRs might possess an extracellular K⁺-sensing mechanism that modulates their ion selectivity in response to ambient K⁺ levels. It has been reported that extracellular K⁺ concentrations in the hippocampal CA3 region of an epileptic mouse model can reach up to 9.2 mM during generalized seizures more than four times the physiological baseline [43]. By analogy, prolonged activation of KCRs could hypothetically result in elevated extracellular K⁺ levels due to K⁺ efflux through the channels, which in turn may lower the P_K_/P_Na_ ratio and compromise the inhibitory effect.

In addition to biophysical factors discussed above, ChR expression levels and subcellular localization may influence the functional outcomes of KCR activation. However, we did not observe significant differences in expression levels or localization patterns among several KCR variants in *Drosophila* motor neurons (**Supplemental Figure 5A-F**). Similarly, in *C. elegans* body wall muscles, the KCR1 and KCR1-C29D expression levels were comparable (**Figure 5D**). The soma-targeted KCR constructs, inspired by the mammalian soma-targeted ACR1 design [17], did not significantly improve inhibitory outcomes in *Drosophila* (**Supplemental Figure 5 B,D**). This highlights the need to identify *Drosophila*-specific subcellular targeting sequences. Targeting light-gated ion channels to distinct neuronal compartments could reveal novel ways to influence action potential generation and neural activity. In addition, differences in temperature, membrane potential, ion homeostasis, and buffering capacity might also influence the extent of KCR-induced hyperpolarization or depolarization.

Collectively, our data emphasizes that both the absolute value and dynamic changes in the P_K_/P_Na_ ratio under illumination must be considered for the effective application of KCRs as inhibitory tools across species. Future engineering of KCRs should focus on rational design approaches that target residues controlling ion permeability shifts and future studies should systematically investigate how environmental factors, such as ionic milieu, temperature, and membrane lipid composition influence the functional properties of KCRs. The optimal KCR will not only be valuable as an inhibitory tool but also offer a deeper understanding of the fundamental mechanisms of neuronal activation and inhibition across organisms, especially when combining different KCRs with subcellular targeting.

## Acknowledgments

We thank Maria Oppmann and Uta Strobel for their support of the *Drosophila* work. We are grateful to Aldrich Hezekiah for providing the initial drawing of adult *Drosophila*.

## Funding

This study was supported by the Deutsche Forschungsgemeinschaft (DFG, German Research Foundation) under project numbers 538090107 and 525167920 awarded to SG, and 426503586 to RJK. SG also acknowledges support from the Homeworld Garden Grants: Protein Engineering for Climate, which contributed to part of the *Xenopus* oocyte experiments. SO, ZZ and ACC were supported by MOE-T2EP30222-0018 grant from the Ministry of Education, Singapore and a Duke-NUS Medical School Block grant. CR is supported by a Kekulé stipend of the Fonds der Chemischen Industrie, and by core funds from Goethe University to AG.

### Author contributions

**XD**: methodology, investigation, formal analysis, data curation, visualization, writing– review and editing. **CZ**: methodology, investigation, formal analysis, data curation, visualization, writing–review and editing. **SO**: project administration, methodology, investigation, formal analysis,data curation, visualization, writing–review and editing. **ZZ**: methodology, investigation, formal analysis, data curation, visualization, writing–review and editing. **CR**: investigation, formal analysis, data curation, visualization, writing– review and editing. **SD**: investigation, formal analysis, visualization, writing–review and editing. **RJ**: investigation, visualization, writing–review and editing. **NE**: investigation, writing–review and editing. **DS**: investigation, writing–review and editing. **GN**: supervision, writing–review and editing. **MH**: supervision, writing–review and editing. **RJK**: initiated the project, supervision, writing–review and editing, and funding acquisition. **AG**: conceptualization, supervision, writing–review and editing, and funding acquisition. **ACC**: initiated the project, project administration, conceptualization, writing– review and editing, supervision, and funding acquisition. **SG**: initiated the project, project administration, conceptualization, methodology, investigation, data curation, visualization, writing–original draft, writing–review and editing, supervision, and funding acquisition.

### Data and materials availability

All data and materials supporting the findings in the manuscript are available from a corresponding author upon reasonable request.

## Methods

### *Xenopus* oocytes: Gene cloning, mRNA preparation, and expression

The KCR1 2.0 construct was previously described [28]. The WiChR DNA was de novo synthesized (GeneArt Strings DNA Fragments, Thermo Fisher Scientific) with additional sequences for BamHI and XhoI (restriction enzymes) at the N-terminal and C-terminal respectively. After enzymatic digestion, the fragment was inserted into the precast pGEMHE vector which contained the cleavable N-terminal signal peptide Lucy-Rho (LR), eYFP, the plasma membrane trafficking signal (T), and the ER export signal (E). For the wild-type version, the fragment was inserted into the precast pGEMHE vector containing the fragment of eYFP. Mutations were obtained via quikchange site-directed mutagenesis. The correct plasmid was linearized by restriction enzyme NheI after the expression cassette. The linearized plasmid was then transcribed in vitro with the AmpliCap-Max™ T7 High Yield Message Maker Kit to obtain mRNA. Thirty nanograms of mRNA were injected into the *Xenopus* oocytes. The oocytes were cultured in ND96 solution containing 10 µM all-*trans* retinal at 16°C. *Xenopus laevis* surgery for oocytes was performed under License #70/14 from Landratsamt Würzburg Veterinaeramt.

### *Xenopus* oocytes: Two-electrode voltage clamp

Photocurrents were measured 2 days after mRNA injection with a light source from an adjustable 530 nm LED (Thorlabs Inc). Electrophysiological measurements with *Xenopus* oocytes were performed at room temperature (∼20°C) with a two-electrode-voltage clamp amplifier (TURBO TEC-05 NPI electronic GmbH, Tamm, Germany). The bath solutions for electrophysiological recordings are indicated in the corresponding figure legends. Electrode capillaries (Ф = 1.5 99 mm, wall thickness 0.178 mm, Hilgenberg) were filled with 3 M KCl, with tip openings of 0.4–1 MΩ. A USB-6221 DAQ device (National Instruments) and WinWCP (v5.5.3, Strathclyde University, United Kingdom) were used for data acquisition. GraphPad Prism 10 software was used for data analysis and Origin2021 Pro software for drawing current traces.

Concentrations of all components are in mM.

**Table 1.** Bath solutions for *Xenopus* oocyte electrophysiology

|  | NaCl | KCl | BaCl<sub>2</sub> | MgCl<sub>2</sub> | HEPES | pH |
| --- | --- | --- | --- | --- | --- | --- |
| Buffer Na | 110 | 0 | 2 | 1 | 5 | 7.4 |
| Buffer high Na | 110 | 5 | 2 | 1 | 5 | 7.4 |
| Buffer High K | 5 | 110 | 2 | 1 | 5 | 7.4 |

### *Drosophila*: Constructs and transgenesis

For transgenic fly generation, the DNA inserts were subcloned from the pGEM vector into the expression vector pJFRC7 using BamHI and HindIII restriction sites. The KALI1 and KALI2 *Drosophila* constructs were obtained by site-directed mutagenesis of the KCR1-ET sequence [18, 22, 26], where the histidine at position 225 was replaced by lysine (Genscript). The synthesized constructs were injected into flies and targeted attP1 or attP2 insertion sites on the second or third chromosomes, respectively. The transgenic progeny was balanced either over CyO or TM6C (BestGene). Expression was verified by imaging of eYFP fluorescence with a Leica TCS SP8 STED confocal microscope.

### *Drosophila* stocks and husbandry

The following *Drosophila* constructs were generated in this study: *y^1^ w^1118^; 20XUAS-KCR1 2.0/CyO*; *y^1^ w^1118^; 20XUAS-WiChR-T95C/CyO*; *y^1^ w^1118^; 20XUAS-sWiChR-T95C/CyO*; *y^1^ w^1118^; 20XUAS-KCR1-TAGA/CyO*; *y^1^ w^1118^; 20XUAS-sKCR1-TAGA/CyO*; *y^1^ w^1118^; 20XUAS-KALI1/CyO*; *y^1^ w^1118^; 20XUAS-KALI2/CyO*. Transgene injections were performed by BestGene Inc. (USA).

The following *Drosophila* stocks were obtained from the Bloomington Drosophila Stock center (BDSC): *elav-GAL4* (BDSC #458), *OK371-GAL4* (BDSC #26160), *MB320C-GAL4* (BDSC #68253), *PPK-GAL4* (BDSC #32078), *Gr5a-GAL4* (BDSC #57591). *UAS-KCR1-ET* and *UAS-KCR1-C29D* stocks have been previously described [18]. The ACR1 stocks have been previously described [29, 30]. The *MB247-GAL4* stock was obtained from Hiromu Tanimoto (Tohoku University, Japan). The *G7-GAL4* stock was requested from Aaron DiAntonio (Washington University, USA). The *OK6-GAL4* stock was obtained from Aberle et al. [44].

Unless otherwise indicated, the *w^1118^* strain was used as a “wild type” control. All *Drosophila* crosses, except for the PER experiments, were prepared on cornmeal-agar medium (1.25% w/v agar, 10.5% w/v dextrose, 10.5% w/v maize, 2.1% w/v yeast) that was supplemented with 0.5 mM all-*trans* retinal. For PER experiments the crosses were prepared on cornmeal-agar without all-trans-retinal and after eclosion the male offspring were transferred to cornmeal-agar supplemented with 0.5 mM all-*trans* retinal. Experimental flies were cultured in vials that were covered in aluminium foil. All *Drosophila* stocks were maintained at 25 °C in temperature-controlled incubators.

### *Drosophila*: Activity monitoring of larvae

The experiments were performed with minor modifications to the previously described protocol [20]. To achieve total internal reflection (TIR) for better image contrast, petri dishes were filled with 1.5% (w/v) agarose, and a 750 nm LED backlight was applied to the lateral surface of the dish. Third instar larvae were collected and transferred onto the agarose surface, where they were allowed to crawl freely. The irradiance lights for optogenetic tools were generated from 450 nm (XXM) or 530 nm (for GtACR1 and KCR1 2.0) LED matrix and powered by a LEDD1B LED Driver (Thorlabs), except that the *PPK>XXM* larvae were illustrated with the light from a 473 nm solid-state laser. The LED Driver was triggered via a 5V signal generated by a relay, and the laser was triggered by a USB-6221 DAQ device (National Instruments). A commercial 5-megapixel CCD camera with 30 FPS was used for video capture. The built-in low-pass 650nm filter was removed, and a long-pass 700 nm filter was added between the sensor and lens.

### *Drosophila*: Posture assessment of larvae

The larvae locomotion assay has been previously described [18]. The 84 × 90 mm assay cassette consisted of 30 discorectangle chambers (26 × 4 mm) that were arranged in two rows. Chambers were cut from 1.5 mm-thick transparent acrylic, coated with 3% agarose and backed with a black sheet of cardboard paper. One third-instar larva was loaded into each chamber in the dark and the chambers were covered with a transparent acrylic lid. The cassette was placed horizontally and illuminated by a mini-projector (Optoma ML750) positioned above the cassette. The larvae were recorded under infrared (IR) light at 24 FPS. Each video frame was processed in real time with CRITTA LabView software [45] and larvae behavioral states in each frame were manually assessed. Activity monitoring experiments consisted of 3 epochs, (1) starting with 45 s of darkness (i.e., infrared illumination only), (2) then followed by 45 s LED illumination, and (3) a final 45 s epoch of darkness. The following illumination parameters were used: red light λ = 617 nm, 13 μW/mm^2^; green light λ = 530 nm, 22 μW/mm^2^; blue light λ = 460 nm, 21 μW/mm^2^. The larvae behavioral states during opsin actuation (**Figure 2A**) are described below: (1) Locomotive = larvae exhibit normal forward crawling behavior. (2) Restricted = larvae display limited movement. (3) Immobile = larvae remain stationary. (4) Contracting = larvae undergo contraction, leading to shortening of body length. (5) Curling = larvae become immobile and adopt a C-shaped posture. (6) Rolling = larvae perform C-shaped body bending and lateral rolling.

### *Drosophila*: Monitoring adult activity

The Trumelan horizontal locomotion tracking assay has been previously described [18]. Briefly, the assay cassette consisted of 26 rectangular chambers arranged in two rows that were cut into an acrylic plate. The back and front of the cassette was covered by a thin sheet of acrylic. In addition, a sheet of black cardboard was placed at the back of the cassette to provide contrast. 1 fly per chamber was loaded into the cassette and the cassette was placed vertically in a Sanyo MIR-154 incubator that was set to 25 °C. *Drosophila* activity was recorded at 10 FPS with a FLIR Grasshopper3 near-infrared video camera (Edmund Optics, GS3-U3-41C6NIR-C) equipped with a 50 mm fixed-focus lens (Edmund Optics, VS-C5024-10M) and an 850 nm long pass filter (Green.L, 58-850). Two sets of infrared LEDs (850 nm peak emission) provided continuous illumination throughout the recording. Each video frame was processed in real time with CRITTA LabView software [45] and fly posture in each frame was manually assessed. Activity monitoring experiments consisted of 3 epochs, (1) beginning with 60 s of darkness (i.e., infrared illumination only) (2) followed by 60 s LED illumination, and (3) a final 60 s epoch of darkness. Illumination was carried out with the following parameters: red light λ = 617 nm, 14 μW/mm^2^; green light λ = 530 nm, 15 μW/mm^2^; blue light λ = 460 nm, 18 μW/mm^2^. The adult fly behavioral states during opsin actuation are described as follows: (1) Supine immobile: flies are positioned upside (2) Supine with leg movement: flies lie upside down while displaying continuous leg movements (3) Curled abdomen: flies adopt a stationary posture with abdominal contraction and curling. (4) Prone: flies are oriented upright but remain stationary. (5) Spastic: flies display a rigid posture and cramped muscles, often progressing from an inclined position to a supine position with leg folding. (6) Restricted walking: flies exhibit limited locomotor activity. (7) Locomotive: flies display normal walking behavior.

### *Drosophila*: Olfactory memory assessment

Aversive olfactory conditioning was performed in a multifly olfactory trainer as previously described [37]. 6 flies were loaded into each chamber and up to 10 chambers were tested simultaneously. Each chamber consisted of a behavioral arena where 3-octanol (OCT) and 4-methylcyclohexanol (MCH) odors were puffed into each chamber end by carrier air adjusted by mass-flow controllers (Sensirion AG). During the 60 s shock-odor training epoch the flies received 12 electric shocks at 60 V that were delivered through the circuit boards on the chamber floor and ceiling. Fly position was recorded at 25 FPS with an AVT F-080 guppy camera that was connected to a video acquisition board (PCI-1409, National Instruments). In each experiment, the flies were conditioned to avoid the shock-paired odor and the post-training odor preference was tested by exposing one half of the chamber to the shock-paired odor and the other half to the unpaired odor. A PI was calculated [46, 47] by counting the position of individual flies in respective video frames over the final 30 s of testing. Conditioned odor avoidance was tested twice - first in the presence and then in the absence of opsin actuation. Shock-odor avoidance during the light on period was compared between genotypic controls and opsin-expressing test flies. Offspring of *w^1118^* flies crossed with the driver stock and offspring of *w^1118^* flies crossed with the UAS-responder stock were used as genotypic controls.

### *Drosophila*: Aversive memory induction

Previous studies have shown that activity in dopaminergic PPL1 neurons induces the formation of aversive odor memory in *Drosophila* [36]. To investigate whether opsin actuation would induce or silence neuronal activity, CsChr, ACR1 and KCRs were expressed in a subset of PPL1 cluster neurons that are captured by the driver line *MB320C-GAL4*. The experiment was performed in the multifly olfactory trainer using a standard conditioning protocol [37], where the shock stimulus was substituted by light-induced opsin actuation during training. Avoidance of the opsin-paired odor was compared against odor bias in the absence of opsin actuation for the respective fly and a mean difference effect size was calculated.

### *Drosophila*: Statistical analysis for adult behavior experiments

Estimation statistics were used to analyze quantitative behavioral data with the DABEST software library [48] as previously described [18]. Mean-difference effect sizes were calculated between the control and the test genotypes. Bootstrap methods were applied to calculate the distributions and 95% CIs of the differences between the tested groups. Data analysis was performed and visualized with Jupyter Python notebooks calling the DABEST, pandas and seaborn packages.

### *Drosophila*: Electrophysiology

*Drosophila melanogaster* 3rd instar larvae of *w^1118^* (control) and *OK6-GAL4/ UAS-KCR1-C29D* strains were used for electrophysiological experiments. Electrophysiological recordings (Axoclamp 2B amplifier, Digidata 1440A; Molecular Devices) were obtained from abdominal muscle 6 in segments A2 and A3 as previously described [49, 50]. All measurements were obtained at room temperature (20 ± 1 °C) in HL-3 [51] with the following composition (in mM): NaCl 70, KCl 5, MgCl_2_ 20, NaHCO_3_ 10, trehalose 5, sucrose 115, Hepes 5, and CaCl_2_ 1, pH adjusted to 7.2. Intracellular electrodes had resistances of 10–30 MΩ and were filled with 3 M KCl. For analysis, only cells with an initial membrane potential of at least −50 mV and a membrane resistance of ≥ 4 MΩ were included. During TEVC recordings, cells were clamped at a holding potential of −60 mV. To evoke synaptic currents, nerves were stimulated via a suction electrode with pulses of 300 μs length and typically at 5-12 V (Grass S48 stimulator and isolation unit SIU5; Astro-Med). For light stimulation, a 530 nm light source (M530F2 – 530 nm, 6,8 mW, ThorLabs; LEDD1B LED Driver, ThorLabs) was triggered via TTL 5V signal (Digidata 1440A, Molecular Devices). Applied light intensity was validated before experiments using a photometer (PLUS2, Laserpoint). Signals were digitized at 10 kHz, low-pass filtered at 1 kHz and analyzed in Clampfit 11.1 (Molecular Devices).

### *Drosophila*: Immunohistochemistry

Stainings were performed essentially as described previously [49]. In brief, following dissection in hemolymph-like solution (HL-3, [51]) CNS attached third instar larval filets were fixed for 10 min in 4% PFA. The samples were then blocked for 30 min in PBT (PBS containing 0.05% Triton X-100, Sigma-Aldrich) and 5% NGS (Dianova) and stained in blocking solution at 4°C overnight. Primary antibodies used were rabbit α-gfp (1:1000, Invitrogen). Samples were washed in PBT and secondary antibody staining was performed in room temperature blocking solution over 3 h. Secondary antibodies used were goat anti-rabbit conjugated Alexa488 (1:500, Invitrogen) and Cy3-conjugated anti-horseradish-peroxidase (anti-hrp, 1:500, Jackson ImmunoResearch). Filets were washed again in PBT and VNCs were removed. The Samples were then mounted in Vectashield (Vector Laboratories, Burlingame, CA, United States) for optical sectioning microscopy. Z-stacks with section thickness 0.3 µm were acquired on an ApoTome system (Zeiss, Jena, Germany, Axiovert 200M Zeiss, objective 63x, NA 1.4, oil). For NMJ imaging, junctions on larval muscles 6 and 7 in segments A2 and A3 were used. For VNC imaging, up to half of the VNC was imaged at the tip where motor neuron somata targeted by the used *OK6-GAL4* driver line should be visible.

### *Drosophila*: Mechanical nociception assay

The experiment was performed based on the protocol from [52] with slight modifications. Third star larvae were poked with a self-made filament at the dorsal A8 segment area. To prepare the filament, a blocked micropipette (Ø 164 um) from World Precision Instruments was cut to a length of 20 mm and attached to a stick with an angle of around 90°. The force of the filament was 60 mN which was calibrated by pressing it to a balance. A sylgard (SylGard 182 Silicone Elastomer Kit; Dow Corning) filled petri dish was prepared for larvae crawling. Before transferring the larva on the sylgard, a ∼1.8 cm^2^ light spot was generated from a fiber. A 650 nm red light from LED was used as a background light. Poke the larva immediately once it was placed on the sylgard.

### *Drosophila*: Proboscis response assay

Prior to the proboscis extension response (PER) experiment, adult flies were collected into a vial with a wet tissue and left for 2 hours. Subsequently, the wings of the adult flies were fixed onto a glass slide using korasilon paste. The fly was carefully adjusted to lie on its side to enable clear visualization of the proboscis by the camera. A 750 nm red light served as background light, and the flies were allowed to remain under this background light for 15 minutes. For illumination, a blue (450 nm) or green (530 nm) light LED matrix was employed. The LED matrix was powered by an LEDD1B LED Driver (ThorLabs), which was triggered by a 5 V signal generated from a relay. A 5-megapixel CCD camera operating at 30 FPS was mounted on a stereo microscope for visual observation and video recording.

In an alternative PER assay setup, the wings of adult flies were fixed onto a glass coverslip using nail polish and the flies were allowed to rest for 30 min in darkness. The coverslip was placed vertically under a Digital Sight 1000 microscope (Nikon) to allow clear visualization of the proboscis by the microscope. Blue (460 nm) or green (530 nm) mobile LEDs were used to illuminate the flies and the light intensity was gradually increased from 0 to 320 μW/mm². The minimum light intensity eliciting a proboscis extension response was recorded for each fly.

### *C. elegans*: Molecular biology and maintenance

The generation of the DNA construct for expression of KCR1 (WT) in body wall muscles of *C. elegans* (*pmyo-3::HcKCR1::mCherry::SL2::GFP*) has been described previously [27]. For expression of KCR1-C29D, a two base pair mutation was introduced through mutagenic PCR using the following primers: 5’-CCCACTTCTTGGATCCATCGACGCTGTTTGCTGCGTTTTC-3’ (forward) and 5’-GAAAACGCAGCAAACAGCGTCGATGGATCCAAGAAGTGGG-3’ (reverse). Transgenic strains were created through microinjection of the DNA constructs into the distal gonads of young adult animals as described previously [53]. The following strains were generated and used in this work: **ZX4083**: N2; *zxEx1578[pmyo-3::HcKCR1::mCherry::SL2::GFP]* (Strain 1), **ZX4084**: N2; *zxEx1579[pmyo-3::HcKCR1::mCherry::SL2::GFP]* (Strain 2), **ZX4085**: N2; *zxEx1580[pmyo-3::HcKCR1(C29D)::mCherry::SL2::GFP]* (Strain 1) and **ZX4086**: N2; *zxEx1581[pmyo-3::HcKCR1(C29D)::mCherry::SL2::GFP]* (Strain 2). All animals were maintained on NGM plates seeded with *Escherichia coli* OP50 bacteria, as described previously [54].

### *C. elegans*: Fluorescence microscopy

Microscopy images were obtained on a Zeiss Axio Observer Z1 using a 40x oil immersion objective. Young adult transgenic animals were immobilized using 200 µM tetramisole hydrochloride (Sigma-Aldrich) on agarose pads made of 7% agarose (Carl Roth) in M9 buffer. Images were processed using FIJI (ImageJ), and partially processed using the “Image Stitching” plugin [55].

### *C. elegans*: Body length assays

Body length assays were performed as described previously [56, 57]. Briefly, on the day prior to the experiments and using red-light filtered transmission light, transgenic L4 larvae were picked onto NGM plates seeded with *E. coli* OP50 bacteria, optionally supplemented with 200 µM all-*trans* retinal (Sigma-Aldrich). For the recordings, individual animals were transferred to unseeded plates and recorded for 30 s using a Canon Powershot G9 connected to a Zeiss Axioscope A1 microscope. The animals were observed for 5s prior to stimulation, followed by 10 s stimulation with light (1 mW/mm^2^) supplied by a 50 mW HBO lamp filtered through a 535/30 bandpass excitation filter (AHF Analysentechnik) and 15 s observation after stimulation. The start of the video acquisition and the stimulation pattern were controlled by an Arduino UNO device running a custom-written script. The videos were analyzed using a WormRuler version 1.3 [58] and the body length was normalized to the mean of seconds 0 to 4. Values below 0.8 and 1.2 excluded, as they are likely to stem from artifacts during background correction. The results were analyzed using Microsoft Excel, OriginPro (OriginLabs) and GraphPad Prism 8.0.2 (GraphPad Software, Boston, US).

## References

1. Piatkevich KD, Boyden ES: Optogenetic control of neural activity: The biophysics of microbial rhodopsins in neuroscience. Q Rev Biophys 2023, 57:e1.

2. Konrad KR, Gao S, Zurbriggen MD, Nagel G: Optogenetic Methods in Plant Biology. Annu Rev Plant Biol 2023, 74:313–339.

3. Govorunova EG, Sineshchekov OA: Channelrhodopsins: From Phototaxis to Optogenetics. Biochemistry-Moscow+ 2023, 88(10):1555–1570.

4. Emiliani V, Entcheva E, Hedrich R, Hegemann P, Konrad KR, Luscher C, Mahn M, Pan ZH, Sims RR, Vierock J, Yizhar O: Optogenetics for light control of biological systems. Nat Rev Methods Primers 2022, 2(1):55.

5. Nagel G, Szellas T, Huhn W, Kateriya S, Adeishvili N, Berthold P, Ollig D, Hegemann P, Bamberg E: Channelrhodopsin-2, a directly light-gated cation-selective membrane channel. P Natl Acad Sci USA 2003, 100(24):13940–13945.

6. Nagel G, Ollig D, Fuhrmann M, Kateriya S, Musti AM, Bamberg E, Hegemann P: Channelrhodopsin-1: a light-gated proton channel in green algae. Science 2002, 296(5577):2395–2398.

7. Wiegert JS, Mahn M, Prigge M, Printz Y, Yizhar O: Silencing Neurons: Tools, Applications, and Experimental Constraints. Neuron 2017, 95(3):504–529.

8. Zhang F, Wang LP, Brauner M, Liewald JF, Kay K, Watzke N, Wood PG, Bamberg E, Nagel G, Gottschalk A, Deisseroth K: Multimodal fast optical interrogation of neural circuitry. Nature 2007, 446(7136):633–639.

9. Gradinaru V, Zhang F, Ramakrishnan C, Mattis J, Prakash R, Diester I, Goshen I, Thompson KR, Deisseroth K: Molecular and cellular approaches for diversifying and extending optogenetics. Cell 2010, 141(1):154–165.

10. Zhang C, Yang S, Flossmann T, Gao S, Witte OW, Nagel G, Holthoff K, Kirmse K: Optimized photo-stimulation of halorhodopsin for long-term neuronal inhibition. BMC Biol 2019, 17(1):95.

11. Oesterhelt D, Stoeckenius W: Functions of a new photoreceptor membrane. P Natl Acad Sci USA 1973, 70(10):2853–2857.

12. Oesterhelt D, Stoeckenius W: Rhodopsin-like protein from the purple membrane of Halobacterium halobium. Nat New Biol 1971, 233(39):149–152.

13. Chow BY, Han X, Dobry AS, Qian X, Chuong AS, Li M, Henninger MA, Belfort GM, Lin Y, Monahan PE, Boyden ES: High-performance genetically targetable optical neural silencing by light-driven proton pumps. Nature 2010, 463(7277):98–102.

14. Mahn M, Prigge M, Ron S, Levy R, Yizhar O: Biophysical constraints of optogenetic inhibition at presynaptic terminals. Nat Neurosci 2016, 19(4):554–556.

15. Govorunova EG, Sineshchekov OA, Janz R, Liu X, Spudich JL: NEUROSCIENCE. Natural light-gated anion channels: A family of microbial rhodopsins for advanced optogenetics. Science 2015, 349(6248):647–650.

16. Zhou Y, Ding M, Gao S, Yu-Strzelczyk J, Krischke M, Duan X, Leide J, Riederer M, Mueller MJ, Hedrich R et al: Optogenetic control of plant growth by a microbial rhodopsin. Nat Plants 2021, 7(2):144–151.

17. Mahn M, Gibor L, Patil P, Cohen-Kashi Malina K, Oring S, Printz Y, Levy R, Lampl I, Yizhar O: High-efficiency optogenetic silencing with soma-targeted anion-conducting channelrhodopsins. Nat Commun 2018, 9(1):4125.

18. Ott S, Xu S, Lee N, Hong I, Anns J, Suresh DD, Zhang Z, Zhang X, Harion R, Ye W et al: Kalium channelrhodopsins effectively inhibit neurons. Nat Commun 2024, 15(1):3480.

19. Cosentino C, Alberio L, Gazzarrini S, Aquila M, Romano E, Cermenati S, Zuccolini P, Petersen J, Beltrame M, Van Etten JL et al: Optogenetics. Engineering of a light-gated potassium channel. Science 2015, 348(6235):707–710.

20. Beck S, Yu-Strzelczyk J, Pauls D, Constantin OM, Gee CE, Ehmann N, Kittel RJ, Nagel G, Gao S: Synthetic Light-Activated Ion Channels for Optogenetic Activation and Inhibition. Front Neurosci 2018, 12:643.

21. Bernal Sierra YA, Rost BR, Pofahl M, Fernandes AM, Kopton RA, Moser S, Holtkamp D, Masala N, Beed P, Tukker JJ et al: Potassium channel-based optogenetic silencing. Nat Commun 2018, 9(1):4611.

22. Govorunova EG, Gou Y, Sineshchekov OA, Li H, Lu X, Wang Y, Brown LS, St-Pierre F, Xue M, Spudich JL: Kalium channelrhodopsins are natural light-gated potassium channels that mediate optogenetic inhibition. Nat Neurosci 2022, 25(7):967–974.

23. Andrews JP, Geng J, Voitiuk K, Elliott MAT, Shin D, Robbins A, Spaeth A, Wang A, Li L, Solis D et al: Multimodal evaluation of network activity and optogenetic interventions in human hippocampal slices. Nat Neurosci 2024, 27(12):2487–2499.

24. Vierock J, Shiewer E, Grimm C, Rozenberg A, Chen IW, Tillert L, Castro Scalise AG, Casini M, Augustin S, Tanese D et al: WiChR, a highly potassium-selective channelrhodopsin for low-light one- and two-photon inhibition of excitable cells. Sci Adv 2022, 8(49):eadd7729.

25. Duan X, Zhang C, Wu Y, Ju J, Xu Z, Li X, Liu Y, Ohdah S, Constantin OM, Pan Y et al: Suppression of epileptic seizures by transcranial activation of K(+)-selective channelrhodopsin. Nat Commun 2025, 16(1):559.

26. Tajima S, Kim YS, Fukuda M, Jo Y, Wang PY, Paggi JM, Inoue M, Byrne EFX, Kishi KE, Nakamura S et al: Structural basis for ion selectivity in potassium-selective channelrhodopsins. Cell 2023, 186(20):4325–4344 e4326.

27. Ruse C, Liewald J, Seidenthal M, Tillert L, Vierock J, Gottschalk A: Potassium-selective channelrhodopsins can exert hyper- or depolarizing effects in excitable cells of *Caenorhabditis elegans*, depending on experimental condition. Genetics 2025:iyaf083.

28. Lin F, Tang R, Zhang C, Scholz N, Nagel G, Gao S: Combining different ion-selective channelrhodopsins to control water flux by light. Pflugers Arch 2023, 475(12):1375–1385.

29. Mohammad F, Stewart JC, Ott S, Chlebikova K, Chua JY, Koh TW, Ho J, Claridge-Chang A: Optogenetic inhibition of behavior with anion channelrhodopsins. Nat Methods 2017, 14(3):271–274.

30. Konig C, Khalili A, Niewalda T, Gao S, Gerber B: An optogenetic analogue of second-order reinforcement in Drosophila. Biol Lett 2019, 15(7):20190084.

31. Scholz N, Guan C, Nieberler M, Grotemeyer A, Maiellaro I, Gao S, Beck S, Pawlak M, Sauer M, Asan E et al: Mechano-dependent signaling by Latrophilin/CIRL quenches cAMP in proprioceptive neurons. Elife 2017, 6.

32. Yang S, Constantin OM, Sachidanandan D, Hofmann H, Kunz TC, Kozjak-Pavlovic V, Oertner TG, Nagel G, Kittel RJ, Gee CE, Gao S: PACmn for improved optogenetic control of intracellular cAMP. BMC Biol 2021, 19(1):227.

33. Dannhäuser S, Lux TJ, Hu C, Selcho M, Chen JTC, Ehmann N, Sachidanandan D, Stopp S, Pauls D, Pawlak M et al: Antinociceptive modulation by the adhesion GPCR CIRL promotes mechanosensory signal discrimination. Elife 2020, 9.

34. Dawydow A, Gueta R, Ljaschenko D, Ullrich S, Hermann M, Ehmann N, Gao S, Fiala A, Langenhan T, Nagel G, Kittel RJ: Channelrhodopsin-2-XXL, a powerful optogenetic tool for low-light applications. P Natl Acad Sci USA 2014, 111(38):13972–13977.

35. Bergs A, Schultheis C, Fischer E, Tsunoda SP, Erbguth K, Husson SJ, Govorunova E, Spudich JL, Nagel G, Gottschalk A, Liewald JF: Rhodopsin optogenetic toolbox v2.0 for light-sensitive excitation and inhibition in Caenorhabditis elegans. PLoS One 2018, 13(2):e0191802.

36. Claridge-Chang A, Roorda RD, Vrontou E, Sjulson L, Li H, Hirsh J, Miesenbock G: Writing memories with light-addressable reinforcement circuitry. Cell 2009, 139(2):405–415.

37. Kan L, Ott S, Joseph B, Park ES, Dai W, Kleiner RE, Claridge-Chang A, Lai EC: A neural m(6)A/Ythdf pathway is required for learning and memory in Drosophila. Nat Commun 2021, 12(1):1458.

38. Tuthill JC, Wilson RI: Parallel Transformation of Tactile Signals in Central Circuits of Drosophila. Cell 2016, 164(5):1046–1059.

39. Medeiros AM, Hobbiss AF, Borges G, Moita M, Mendes CS: Mechanosensory bristles mediate avoidance behavior by triggering sustained local motor activity in Drosophila melanogaster. Curr Biol 2024, 34(13).

40. de Belle JS, Heisenberg M: Associative odor learning in Drosophila abolished by chemical ablation of mushroom bodies. Science 1994, 263(5147):692–695.

41. Akuzawa-Tateyama M, Tateyama M, Ochi R: Low K+-induced hyperpolarizations trigger transient depolarizations and action potentials in rabbit ventricular myocytes. J Physiol-London 1998, 513 (Pt 3)(Pt 3):775–786.

42. Rama S, Zbili M, Bialowas A, Fronzaroli-Molinieres L, Ankri N, Carlier E, Marra V, Debanne D: Presynaptic hyperpolarization induces a fast analogue modulation of spike-evoked transmission mediated by axonal sodium channels. Nat Commun 2015, 6:10163.

43. Liu J, Li F, Wang Y, Pan L, Lin P, Zhang B, Zheng Y, Xu Y, Liao H, Ko G et al: A sensitive and specific nanosensor for monitoring extracellular potassium levels in the brain. Nat Nanotechnol 2020, 15(4):321–330.

44. Aberle H, Haghighi AP, Fetter RD, McCabe BD, Magalhaes TR, Goodman CS: encodes a BMP type II receptor that regulates synaptic growth in. Neuron 2002, 33(4):545–558.

45. Krishnan S, Mathuru AS, Kibat C, Rahman M, Lupton CE, Stewart J, Claridge-Chang A, Yen SC, Jesuthasan S: The right dorsal habenula limits attraction to an odor in zebrafish. Curr Biol 2014, 24(11):1167–1175.

46. Tully T, Quinn WG: Classical conditioning and retention in normal and mutant Drosophila melanogaster. J Comp Physiol A 1985, 157(2):263–277.

47. Tumkaya T, Ott S, Claridge-Chang A: A systematic review of Drosophila short-term-memory genetics: Meta-analysis reveals robust reproducibility. Neurosci Biobehav R 2018, 95:361–382.

48. Ho J, Tumkaya T, Aryal S, Choi H, Claridge-Chang A: Moving beyond P values: data analysis with estimation graphics. Nat Methods 2019, 16(7):565–566.

49. Dannhauser S, Mrestani A, Gundelach F, Pauli M, Komma F, Kollmannsberger P, Sauer M, Heckmann M, Paul MM: Endogenous tagging of Unc-13 reveals nanoscale reorganization at active zones during presynaptic homeostatic potentiation. Front Cell Neurosci 2022, 16:1074304.

50. Beckers CJ, Mrestani A, Komma F, Dannhauser S: Versatile Endogenous Editing of GluRIIA in Drosophila melanogaster. Cells-Basel 2024, 13(4).

51. Stewart BA, Atwood HL, Renger JJ, Wang J, Wu CF: Improved stability of Drosophila larval neuromuscular preparations in haemolymph-like physiological solutions. J Comp Physiol A 1994, 175(2):179–191.

52. Zhong L, Hwang RY, Tracey WD: Pickpocket is a DEG/ENaC protein required for mechanical nociception in Drosophila larvae. Curr Biol 2010, 20(5):429–434.

53. Fire A: Integrative transformation of Caenorhabditis elegans. Embo J 1986, 5(10):2673–2680.

54. Brenner S: The genetics of Caenorhabditis elegans. Genetics 1974, 77(1):71–94.

55. Preibisch S, Saalfeld S, Tomancak P: Globally optimal stitching of tiled 3D microscopic image acquisitions. Bioinformatics 2009, 25(11):1463–1465.

56. Nagel G, Brauner M, Liewald JF, Adeishvili N, Bamberg E, Gottschalk A: Light activation of channelrhodopsin-2 in excitable cells of Caenorhabditis elegans triggers rapid behavioral responses. Curr Biol 2005, 15(24):2279–2284.

57. Liewald JF, Brauner M, Stephens GJ, Bouhours M, Schultheis C, Zhen M, Gottschalk A: Optogenetic analysis of synaptic function. Nat Methods 2008, 5(10):895–902.

58. Seidenthal M, Vettkotter D, Gottschalk A: WormRuler: A software to track body length used to characterize a super red-shifted channelrhodopsin in Caenorhabditis elegans. MicroPubl Biol 2022, 2022.

