## Supplemental Figures and Tables for "Stabilized ion selectivity corrects activation drift in kalium channelrhodopsins"

### Ion selectivity shifts underlie the dual function of kalium channelrhodopsins in *Drosophila* and *C. elegans*

#### Supplemental Figures

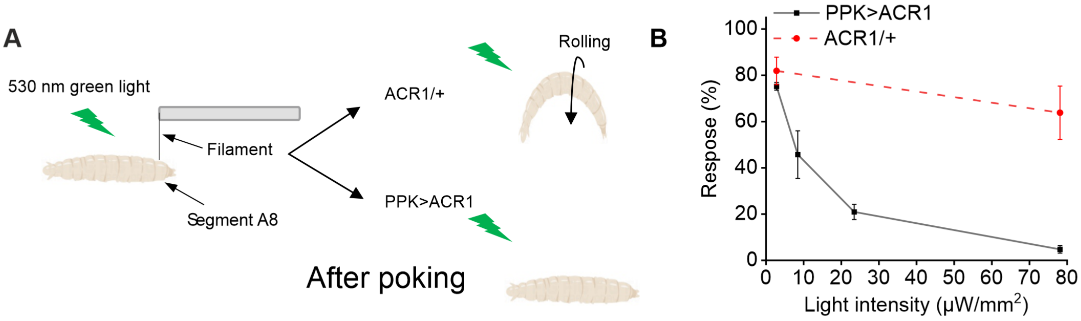


**Supplemental Figure 1: ACR1 efficiently suppresses nociceptive responses in *Drosophila* larvae. A**, Schematic of the experimental setup used to assess antinocifensive effects in response to physical pain stimuli under light illumination. **B**, Proportion of larvae exhibiting corkscrew-like behaviour at different green light (530 nm) intensities. A total of 35 larvae were tested per condition, with n = 3 independent replicates. Data are presented as mean ± SD.

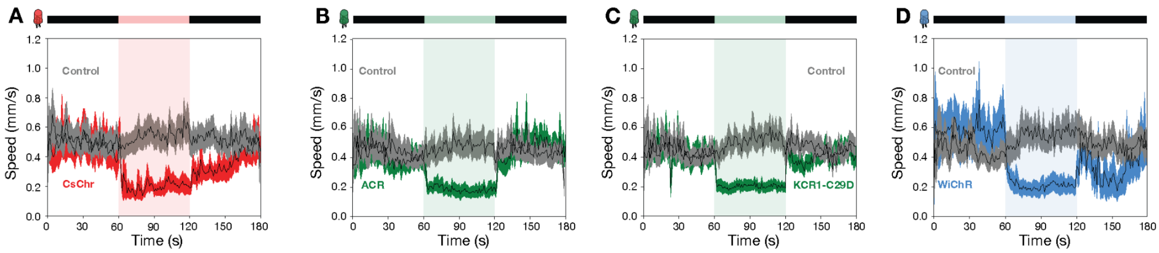


**Supplemental Figure 2: Neuronal activation and silencing reduces *Drosophila* larvae crawling speed.** The panels illustrate *Drosophila* larvae crawling speed before, during and after opsin actuation. ChRs were expressed in motor neurons by using *OK371-GAL4*. Neuronal activation with the cation-conducting channel CsChr (A) as well as actuation with ACR and KCRs (B-D) impaired locomotion. The light illumination period is indicated in the schematic on top of each panel. Genotypic driver and responder controls were used in each experiment. Sample size: *OK371>CsChr* n = 29 larvae, controls n = 58 larvae; *OK371>ACR* n = 30 larvae, controls n = 58 larvae; *OK371>KCR1-C29D* n = 30 larvae, controls n = 58 larvae; *OK371>WiChR* n = 30 larvae, controls n = 58 larvae. Error bands show 95% Confidence interval (CI).

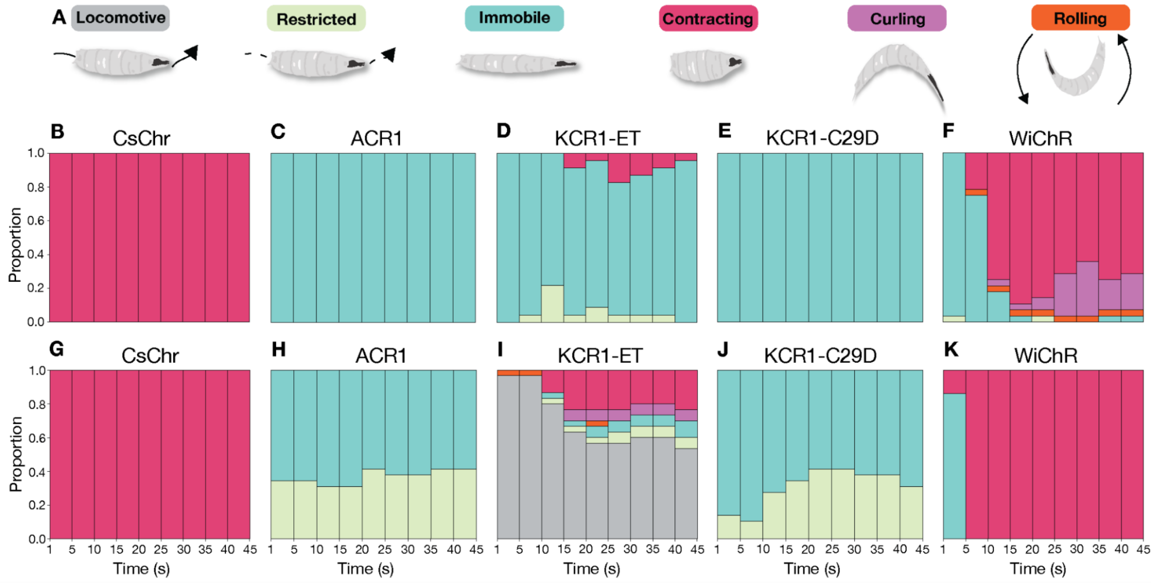


**Supplemental Figure 3: Pan-neuronal and motor neuron KCR actuation elicits different behavioral states. A**, Schematic and colour legend of behavioral states in *Drosophila* larvae induced by opsin actuation. **B-K**, State bar charts of *Drosophila* larvae expressing different opsins pan-neuronally with *elav-GAL4* (**B-F**) or in motor neurons with *OK6-GAL4* (**G-I**). Sample size: *elav>CsChr* n = 30, *elav>ACR1* n = 30, *elav>KCR1-ET* n = 23, *elav>KCR1-C29D* n = 30, *elav>WiChR* n = 28, *OK6>CsChr* n = 30, *OK6>ACR1* n = 29, *OK6>KCR1-ET* n = 30, *OK6>KCR1-C29D* n = 29, *OK6>WiChR* n = 29.

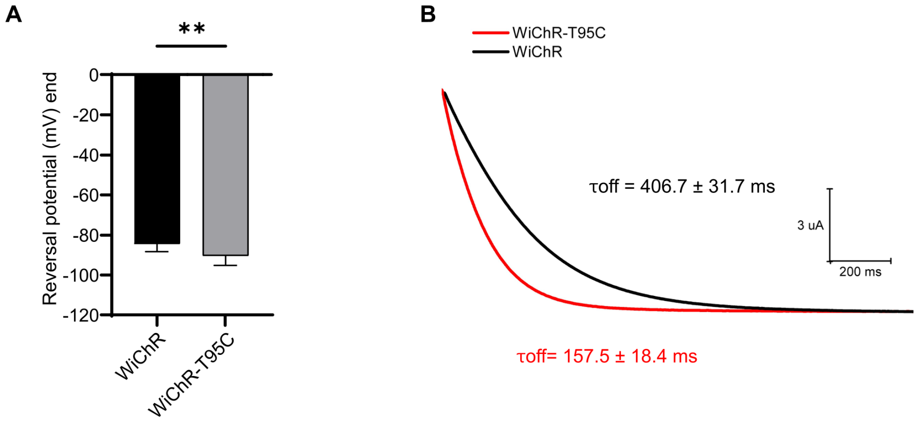

**Supplemental Figure 4: Characterization of the WiChR T95C mutant. A**, Comparison of the reversal potential of WiChR and the T95C mutant in buffer containing 110 mM NaCl. n = 10. **B**, Comparison of closing kinetics after light stimulation. Data are presented as mean ± SD.

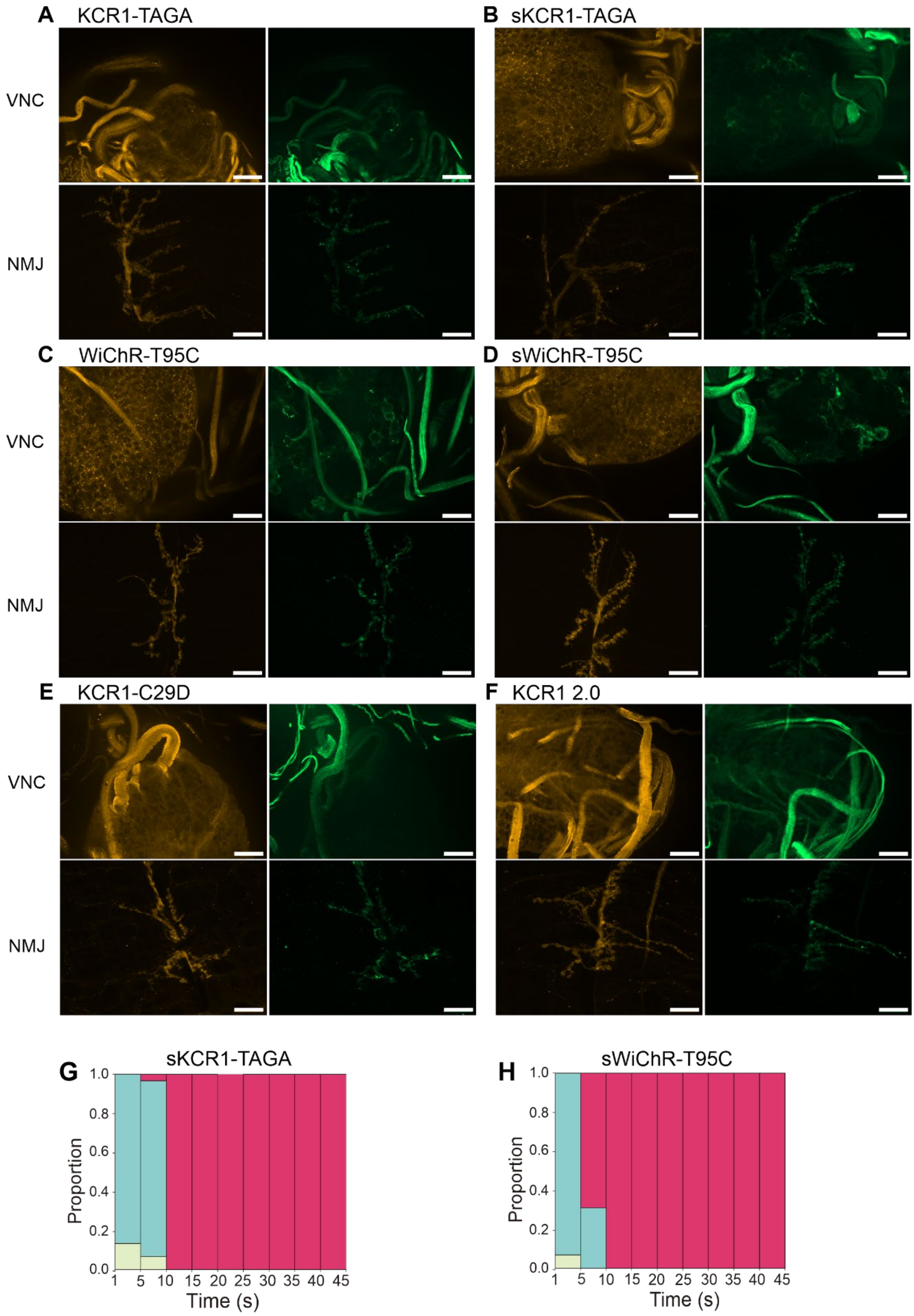


**Supplemental Figure 5: Expression patterns and light-induced behavioral responses of soma-targeted KCRs and other variants in *Drosophila* motor neurons.** Representative maximum intensity Z-projections of VNCs and NMJs of larvae expressing different *OK6-GAL4* driven, YFP-tagged opsins (Hrp: yellow, YFP: green). Images of **(A)** KCR1-TAGA and **(B)** sKCR1-TAGA, **(C)** WiChR-T95C and **(D)** WiChR-T95C as well as **(E)** KCR1-C29D and **(F)**

KCR1 2.0 show the VNC at the top and NMJ at the bottom. Scale bars: 20 µm. **(G)** and **(H)** State bar chart during light illumination for *Drosophila* larvae expressing *OK371>sKCR1-TAGA* and *OK371>sWiChR-T95C*. The illumination epoch (45s) was divided into 5s bins and the prevalence of each state across animals was plotted as a proportion in each bin (see methods).

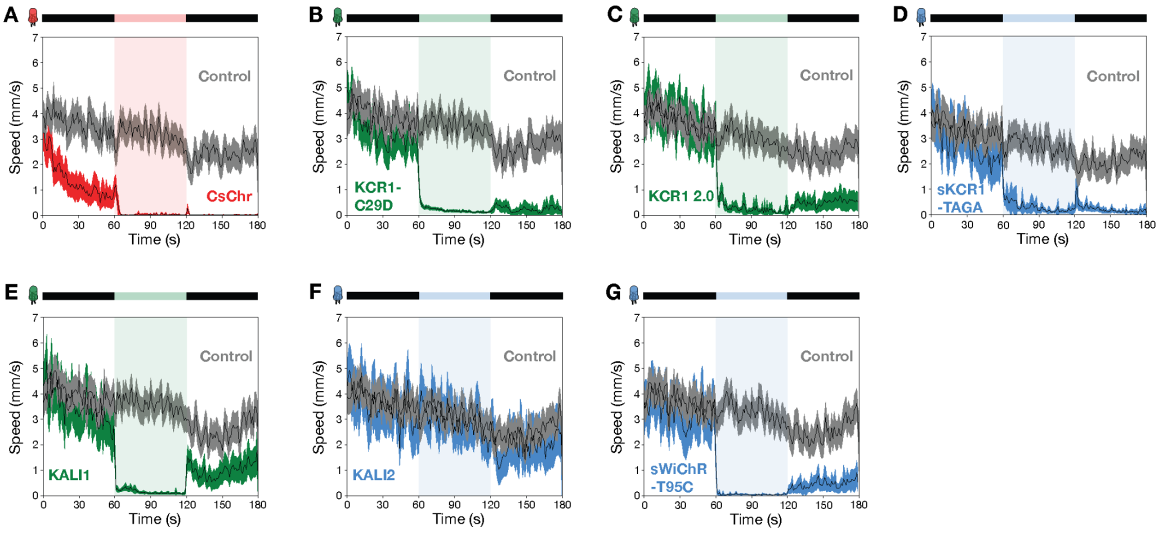


**Supplemental Figure 6: Optogenetic activation and silencing of motor neurons impairs *Drosophila* locomotion.** The panels illustrate adult *Drosophila* locomotor activity before, during and after opsin actuation. Apart from KALI2, the activation of different KCR variants as well as CsChr strongly impaired locomotion. The same data as shown in **Figure 2** was used to analyse *Drosophila* locomotor activity. Sample size: *OK371>CsChr* n = 52, controls = 104; *OK371>KCR1-C29D* n = 52, controls = 104; *OK371>KCR1 2.0* n = 40, controls = 104; *OK371>sKCR1-TAGA* n = 26, controls = 104; *OK371>KALI1* n = 40, controls = 104; *OK371>KALI2* n = 40, controls = 104; *OK371>sWiChR-T95C* n = 26, controls = 104. Error bands show 95% CI.

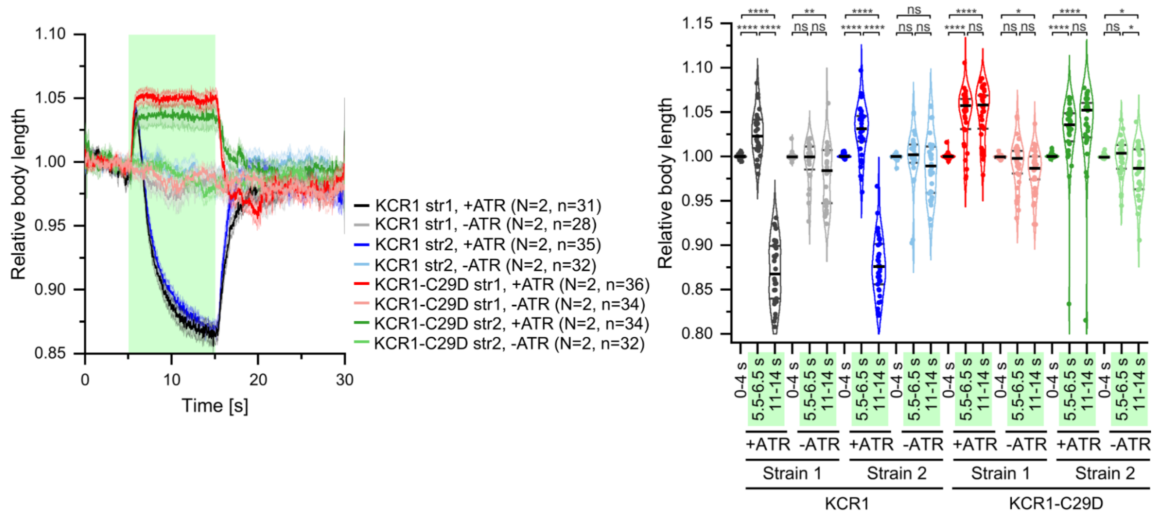

**Supplemental Figure 7:** Time and light-dependent body length profiles of two additional independent *C. elegans* transgenic strains ("str2") expressing KCR1 (WT) or KCR1-C29D in body wall muscles (*pmyo-3::KCR1(WT or C29D)::mCherry::SL2::GFP*, 15ng/µL), compared to the strains ("str1") shown in Figure 5E. The worms were illuminated for 10 s with 535 nm light at an intensity of 1 mW/mm$^2$, with and without additional all-*trans* retinal supplementation. Statistical significance was assessed using a mixed-effect analysis model (REML) with Tukey's multiple comparisons test.

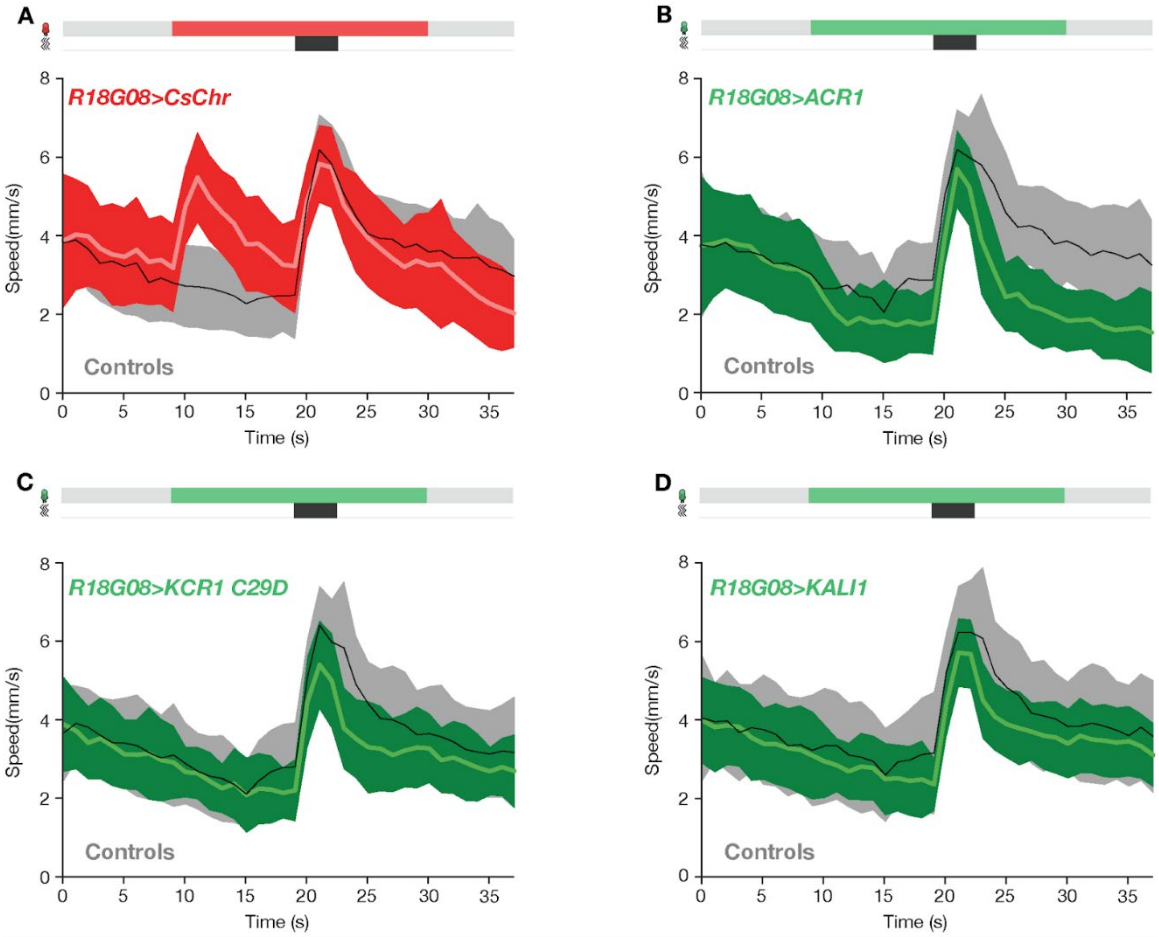


**Supplemental Figure 8: KCR1 C29D and KALI effectively silence *Drosophila* somatosensory neurons.** The panels display the locomotor activity of *Drosophila* before and during somatosensory neuron actuation. Opsins were expressed in second order somatosensory neurons by using *R18G08-GAL4*. The schematic on top of each panel indicates the period of illumination (colored square) and air puff agitation (black square). Activation of somatosensory neurons by CsChr induced locomotor activity (**A**), whereas their silencing by ACR (**B**) and KCRs (**C-D**) impaired the post-agitation locomotor response. Light intensities: red: 24 µW/mm$^2$; green = 58 µW/mm$^2$. Sample size: *R18G08>CsChr* n = 348 flies, controls n = 330 flies; *R18G08>ACR1* n = 348 flies, controls n = 348 flies; *R18G08>KCR1-C29D* n = 354 flies, controls n = 336 flies; *R18G08>KALI1* n = 342 flies, controls n = 348 flies. Error bands show 95% CI.

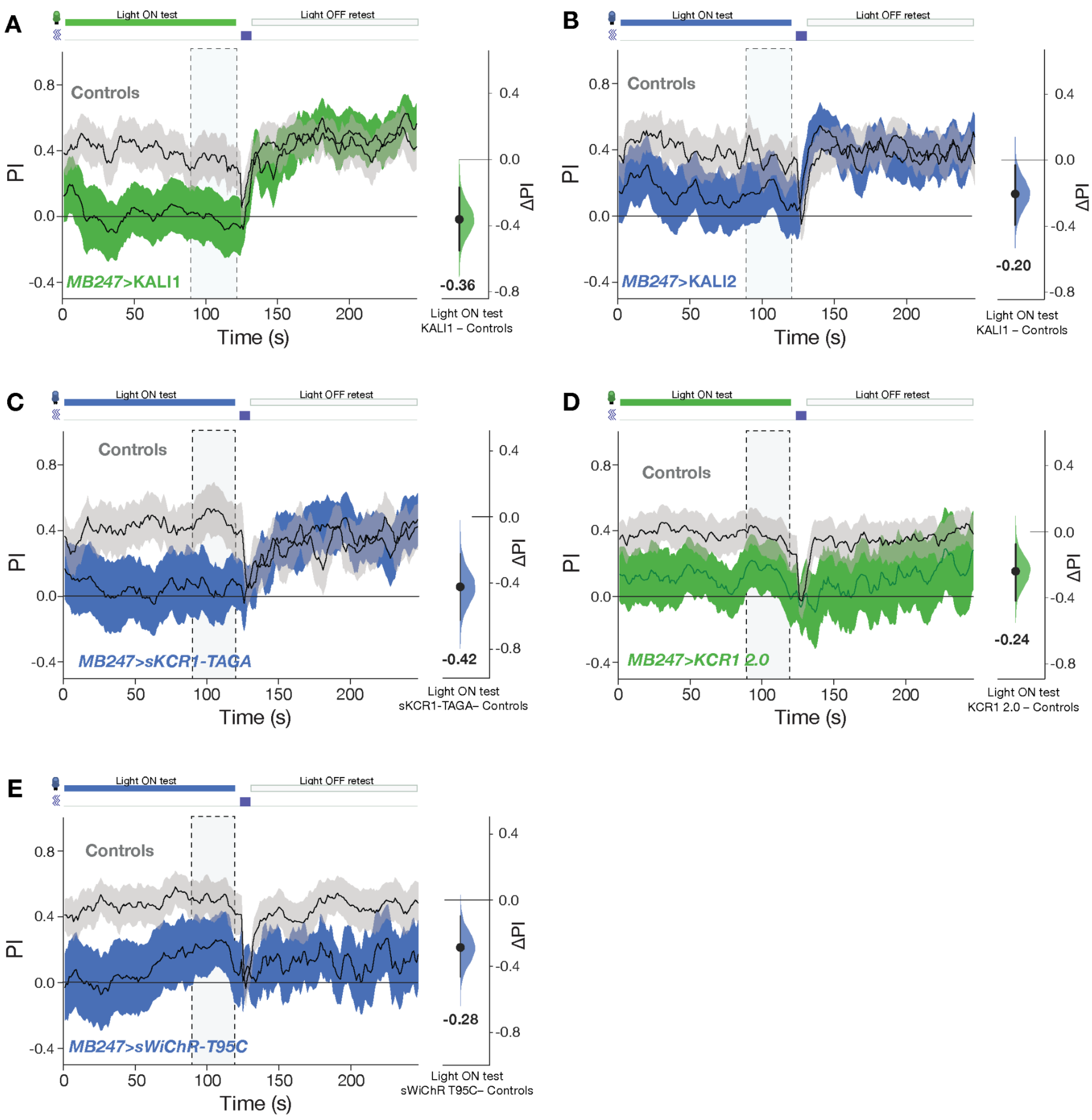


**Supplemental Figure 9: KCR actuation inhibits associative memory. A-E**, The panels show the shock-odor avoidance PI during the light-on and light-off testing epochs. The schematic above shows the period of illumination (filled rectangle), light off retest (empty rectangle) and agitation (blue rectangle). The axis on the right shows the mean difference effect size comparison between the PI of the genotypic driver and responder controls and the test flies. The dashed rectangle indicates the time interval used for effect size comparisons. The silencing of mushroom body neurons with *MB247>KALI1* (**A**) and *MB247>sKCR1-TAGA* (**C**) strongly impaired aversive odor memory and re-testing of the same animals in the absence of light stimulation restored conditioned odor avoidance. MB neuron silencing with KALI2 (B), KCR1 2.0 (D) and sWiChR-T95C (**E**) was less effective. Error bands in all panels represent the 95% CI. Light intensities for all panels: red: 24 μW/mm$^2$; green = 58 μW/mm$^2$, blue 21 μW/mm$^2$. Sample size: *MB247>KALI1* n = 336 flies, controls n = 396 flies; *MB247>KALI2* = 294 flies, controls n = 414 flies; *MB247>sKCR1-TAGA* = 240 flies, controls n = 306 flies; *MB247>KCR1 2.0* = 240 flies, controls n = 480 flies; *MB247C>sWiChR-T95C* = 240 flies, controls n = 480 flies.

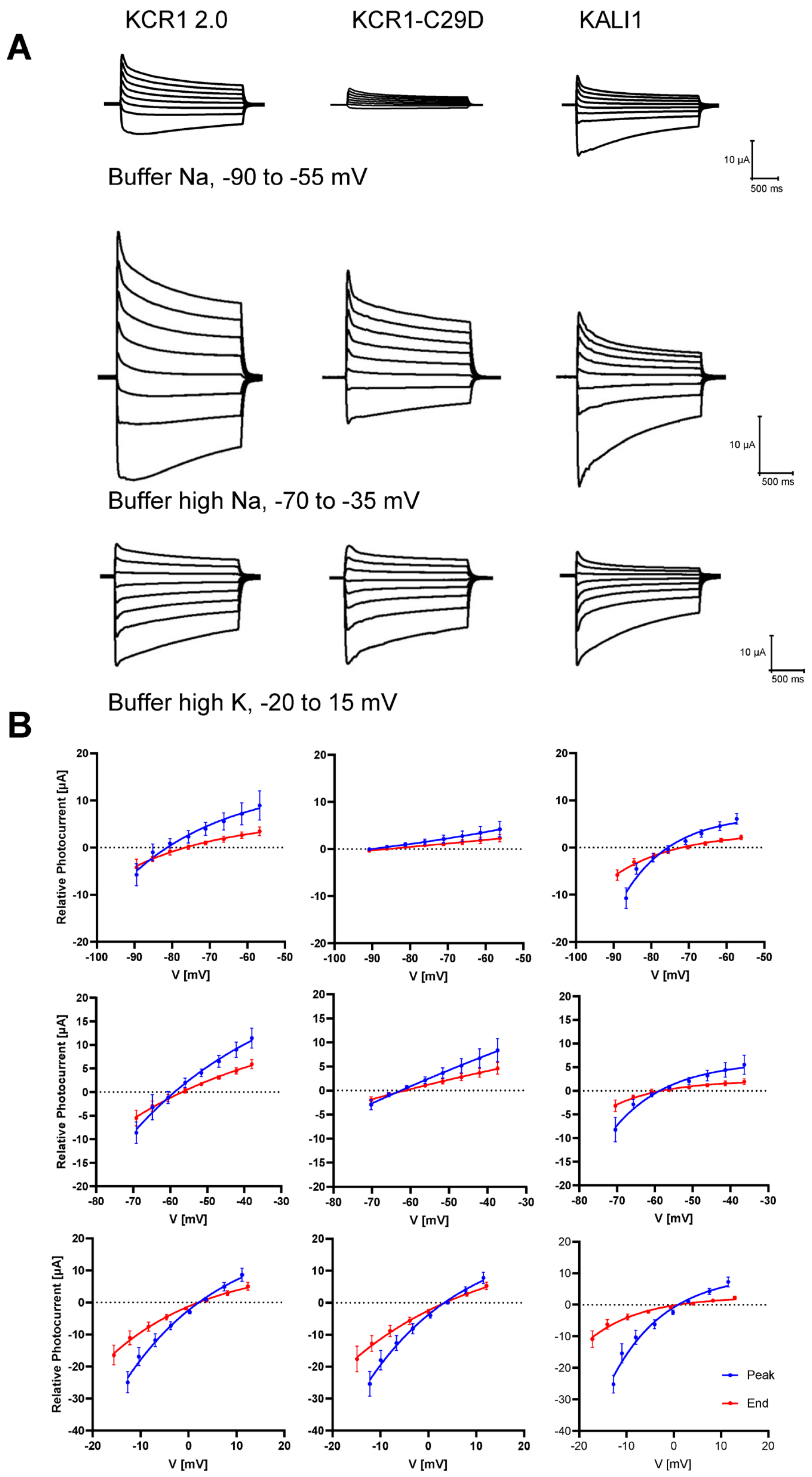


**Supplemental Figure 10: Representative photocurrent traces (A) and I-V curve (B) of KCR1 2.0, KCR1-C29D, and KALI1.** **A**, Representative photocurrent traces were recorded in response to 2-s light pulses under buffer Na, buffer high Na, and buffer high K conditions. Holding voltages were incremented in 5 mV steps. **B**, The corresponding current-voltage (I-V) relationships were plotted, with the blue curve representing peak photocurrent and the red curve representing end photocurrent (mean ± sem, n = 6 - 10).

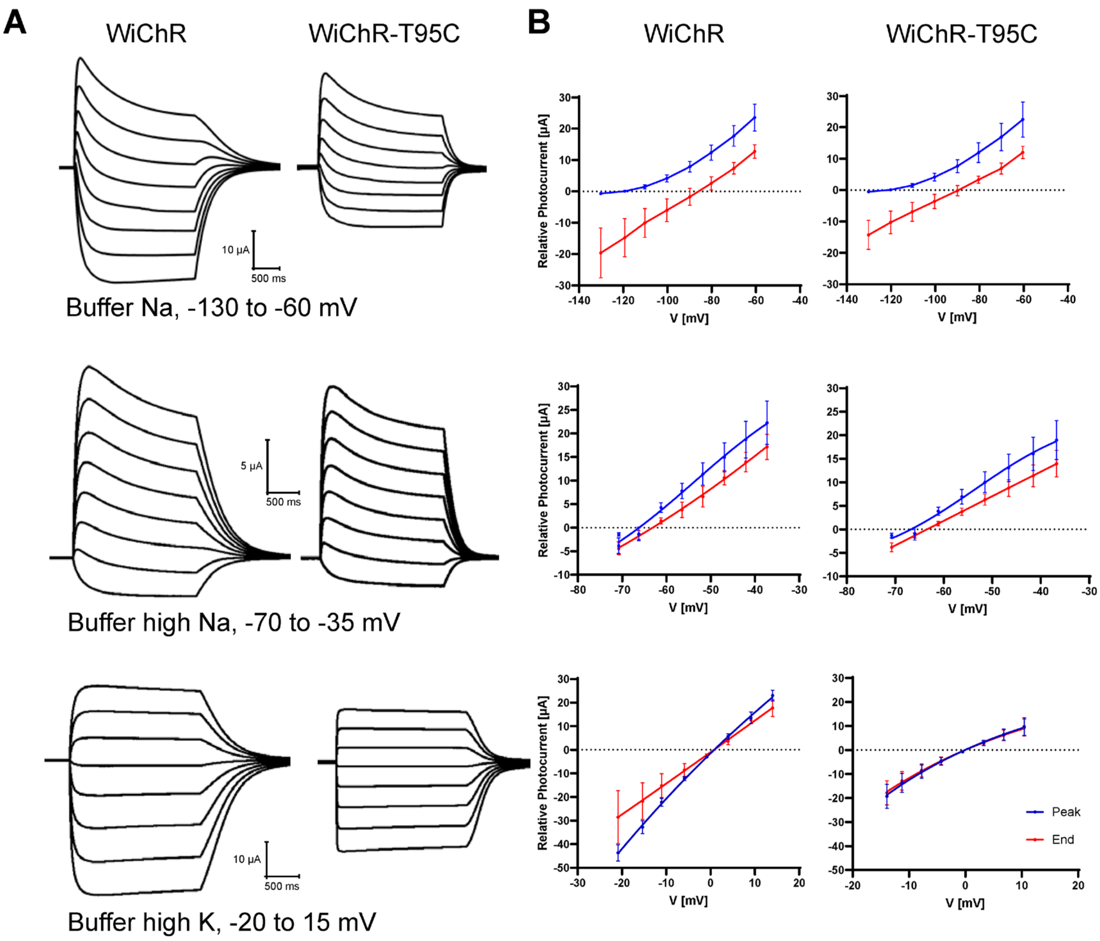


**Supplemental Figure 11: Representative photocurrent traces (A) and I-V curve (B) of WiChR and WiChR-T95C. A**, Representative photocurrent traces were recorded in response to 2 s light pulses under buffer Na, buffer high Na, and buffer high K conditions. Holding voltages were incremented in 10 mV steps under buffer Na and 5-mV steps under buffer high Na, and buffer high K conditions. **B**, The corresponding current-voltage (I-V) relationships were plotted, with the blue curve representing peak photocurrent and the red curve representing end photocurrent (mean ± sem, n = 6 - 10).

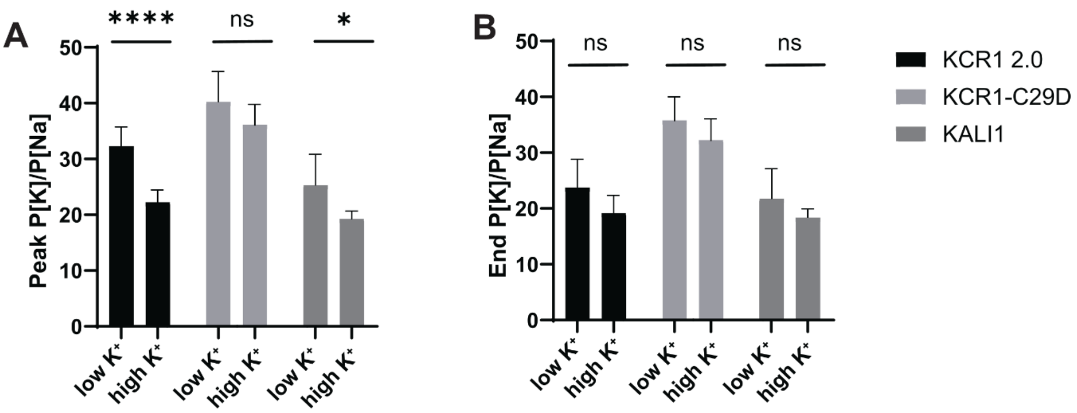


**Supplemental Figure 12: $P_K/P_{Na}$ ratios calculated from different buffer combinations. A**, The permeability ratios calculated using reversal potentials measured at the peak response. low $K^+$: calculations performed according to results from buffer Na and high Na; high $K^+$: calculations performed according to results from buffer high Na and high K. Buffer details in Figure 6A. **B**, The permeability ratios calculated using reversal potentials measured at the end of a 2 s light pulse.

#### Supplemental Tables

##### Supplemental Table 1: Electrophysiological analysis of control and mutant NMJs.

Related to **Figure 3D**. Data are presented in -nA; SD denotes standard deviation; SEM denotes standard error of mean; n denotes number of measured NMJs.

| Genotype light condition | eEPSC amplitude (-nA) | | | | | | n |
|---|---|---|---|---|---|---|---|
| | Minimum | Maximum | Median | Mean | SD | SEM | |
| control ($w^{1118}$) before light 4 µW | 35,20 | 69,65 | 53,89 | 53,90 | 13,04 | 4,61 | 8 |
| control ($w^{1118}$) before light 20 µW | 31,54 | 66,11 | 47,25 | 47,78 | 11,14 | 4,21 | 7 |
| ok6 > $KCR^{C29D}$ before light 4 µW | 53,13 | 71,92 | 61,77 | 61,84 | 8,42 | 2,66 | 10 |
| ok6 > $KCR^{C29D}$ before light 20 µW | 53,86 | 82,40 | 64,33 | 64,48 | 9,37 | 2,71 | 12 |
| control ($w^{1118}$) during light 4 µW | 32,59 | 68,04 | 48,81 | 49,15 | 12,76 | 4,51 | 8 |
| control ($w^{1118}$) during light 20 µW | 27,34 | 58,87 | 42,42 | 43,25 | 9,30 | 3,51 | 7 |
| ok6 > $KCR^{C29D}$ during light 4 µW | -6,25 | 26,66 | 5,62 | 7,69 | 11,17 | 3,53 | 10 |
| ok6 > $KCR^{C29D}$ during light 20 µW | 0,55 | 1,14 | 0,70 | 0,75 | 0,18 | 0,05 | 12 |
| control ($w^{1118}$) after light 4 µW | 29,78 | 71,16 | 48,17 | 49,78 | 15,98 | 5,65 | 8 |
| control ($w^{1118}$) after light 20 µW | 28,96 | 64,80 | 42,22 | 43,97 | 10,65 | 4,03 | 7 |
| ok6 > $KCR^{C29D}$ after light 4 µW | 51,47 | 75,46 | 60,09 | 61,42 | 8,21 | 2,60 | 10 |
| ok6 > $KCR^{C29D}$ after light 20 µW | 21,82 | 94,41 | 60,10 | 59,46 | 19,55 | 5,64 | 12 |

##### Supplemental Table 2: Statistical information related to Table 1 (TEVC recordings).

Kruskal-Wallis test with follow-up Dunn's multiple comparisons test was used. *n.s.* denotes $p > 0.9999$

| **Genotype; light condition** | control; before; 4 µW | control; before; 20 µW | $KCR^{C29D}$; before; 4 µW | $KCR^{C29D}$; before; 20 µW | control; during; 4 µW | control; during; 20 µW | $KCR^{C29D}$; during; 4 µW | $KCR^{C29D}$; during; 20 µW | control; after; 4 µW | control; after; 20 µW | $KCR^{C29D}$; after; 4 µW | $KCR^{C29D}$; after; 20 µW |
|---|---|---|---|---|---|---|---|---|---|---|---|---|
| control; before; 4 µW | / | | | | | | | | | | | |
| control; before; 20 µW | n.s. | / | | | | | | | | | | |
| $KCR^{C29D}$; before; 4 µW | n.s. | n.s. | / | | | | | | | | | |
| $KCR^{C29D}$; before; 20 µW | n.s. | n.s. | n.s. | / | | | | | | | | |
| control; during; 4 µW | n.s. | n.s. | n.s. | n.s. | / | | | | | | | |
| control; during; 20 µW | n.s. | n.s. | n.s. | .4596 | n.s. | / | | | | | | |
| $KCR^{C29D}$; during; 4 µW | .0536 | .9675 | .0002 | < .0001 | .4781 | n.s. | / | | | | | |
| $KCR^{C29D}$; during; 20 µW | .0101 | .3140 | < .0001 | < .0001 | .1290 | n.s. | n.s. | / | | | | |
| control; after; 4 µW | n.s. | n.s. | n.s. | n.s. | n.s. | n.s. | .2172 | .0513 | / | | | |
| control; after; 20 µW | n.s. | n.s. | n.s. | .5737 | n.s. | n.s. | n.s. | n.s. | n.s. | / | | |
| $KCR^{C29D}$; after; 4 µW | n.s. | n.s. | n.s. | n.s. | n.s. | n.s. | .0004 | < .0001 | n.s. | n.s. | / | |
| $KCR^{C29D}$; after; 20 µW | n.s. | n.s. | n.s. | n.s. | n.s. | n.s. | .0004 | < .0001 | n.s. | n.s. | n.s. | / |
